# Organobodies: A robust and size-controllable system for generating scalable hiPSC-derived liver organoids for drug toxicity screening

**DOI:** 10.1101/2025.10.20.683433

**Authors:** Mostafa Kiamehr, Stefano Manzini, Burak Toprakhisar, Rodrigo F. Madeiro da Costa, Guillem García-Llorens, Birhanu Belay, Mustapha Najimi, José V Castell, Wolfgang Moritz, Giulia Chiesa, Katriina Aalto-Setälä, Catherine Verfaillie

## Abstract

**Background and aim:** Hepatic organoids generated from pluripotent and adult stem cells faithfully mimic the architecture and function of *in vivo* organs making them powerful tools for preclinical applications such as disease modelling and drug screening. However, developing robust and scalable models capable of preserving liver-specific functions long-term remains technically challenging.

**Methods:** Using a defined self-assembling peptide, we established a robust and reproducible method to generate hepatic organoids from human induced pluripotent stem cells (hiPSCs), termed hepatic *organobodies* (OBs), which remained stable in culture for several weeks. The hepatic identity and maturation of OBs were characterized by qPCR, histology, immunofluorescence, and bulk RNA sequencing. Their functional competence was evaluated by ELISA, *CYP3A4* induction assays, and assessment of drug biotransformation capacity using HPLC–MS/MS, in parallel with three-dimensional human primary hepatocyte (3D PHH) cultures. Finally, the model was applied to predict the hepatotoxicity of a selected panel of known compounds and benchmarked against HepG2 cells and 3D PHH microtissues.

**Results:** hiPSC-derived hepatocyte-like cells (HLCs) cultured in OBs acquired characteristic hepatic morphology and expressed several key hepatocyte-specific genes, some at levels comparable to those in freshly isolated primary human hepatocytes (PHHs). OBs secreted significantly higher amounts of albumin and α1-antitrypsin (A1AT) compared with parallel 2D-cultured HLCs. Bulk RNA sequencing revealed higher relative expression of major drug metabolism genes, including *CYP3A4*, *CYP2C9*, and *CYP1A2*, as well as enhanced maturation, evidenced by upregulation of the PPAR signalling and fatty acid β-oxidation pathways in OBs relative to 2D HLCs. Moreover, OBs demonstrated CYP450 enzyme activity, with CYP3A4 and CYP2C9 activities comparable to those observed in 3D PHH microtissues. Finally, we show that our 3D model accurately predicted the hepatotoxicity of more than 10 tested chemical compounds.

**Conclusion:** Here, we demonstrate a novel, defined, robust and remarkably advanced 3D liver model as a valuable platform for scalable toxicity prediction and drug screening for personalised medicine.

## Introduction

The liver, as the body’s primary detoxification organ, plays a central role in metabolizing and eliminating xenobiotics. Consequently, more than 20% of newly approved drugs that are later withdrawn from the market are associated with liver injury and hepatotoxicity ^1,2^, a rate that could be significantly reduced through the use of more predictive preclinical models. Traditionally, human health risk assessment has relied on animal studies; however, species-specific differences in anatomy, physiology, and metabolism limit their ability to accurately predict human toxicity ^3,4^. Notably, nearly half of drug candidates that cause hepatotoxicity in humans show no such effect in animal models ^5^. In response, a 2007 report by the U.S. National Research Council called for a transition toward human-relevant alternatives in toxicology, and the U.S. Environmental Protection Agency has committed to phasing out animal testing for chemicals and pesticides by 2035 ^6^. These developments underscore the urgent need for physiologically relevant, human-derived liver models to improve the predictive power of hepatotoxicity screening.

While conventional two dimensional (2D) *in vitro* models facilitate preliminary screening, they lack the physiological complexity of human organs. In contrast, three dimensional (3D) liver models and organoids more closely recapitulate native tissue architecture and are increasingly used for developmental studies, disease modelling, personalized medicine, tissue engineering, drug screening, and adverse drug reaction (ADR) detection ^7–10^.

Several strategies have been developed to generate scalable 3D liver models such as spheroids and organoids. Conventional approaches involve self-aggregation of cells in microwells ^11–13^, nanopillar ^14^ or ultra-low attachment plates ^15^. However, spheroids typically range from 100 to 200 µm in diameter, yield limited amounts of RNA and protein from individual spheroids, and pose handling challenges. Larger spheroids are prone to central necrosis, and multiple spheroids cultured in the same well or chamber often fuse rapidly, increasing variability in size and survival. The hanging drop method, though an alternative, is time-consuming and labour-intensive, particularly when applied to large-scale studies ^16^.

Pluripotent stem cell (PSC)-derived organoids represent a promising and powerful approach; however, their formation typically relies on Matrigel, a complex and poorly defined animal-derived matrix with batch variability and limited clinical relevance ^17^. Additionally, the culture of tissue-derived organoids typically requires routine passaging, restricting experiments to a 1–2-week window and limiting repeated-dose toxicity testing ^4,18,19^.

Recent studies highlight the promising potential of self-assembling peptide (SAP) nanofibers for 3D cell culture, biomedical, and clinical applications due to their excellent bioactivity and biodegradability ^20^. These nanofibers are composed of natural amino acids and form hydrogels with properties that mimic native nanofibrous extracellular matrix (ECM). A well-characterized example is RADA16 (Ac-(RADA)₄-CONH₂), an amphiphilic 16-residue peptide composed of alternating hydrophobic and hydrophilic amino acids. Under neutral pH and physiological salt conditions, such as those found in cell culture media, RADA16 forms stable β-sheets that self-assemble into nanofibers which further organize into a hydrogel scaffold composed of ∼99% water. The resulting nanofibers are approximately 6–10 nm in diameter and organize into a porous network with pore sizes ranging from 5 to 200 nm ^21,22^. RADA16-based scaffolds have been demonstrated to support culture and differentiation of many cell types such as hepatocytes ^23,24^, neurons ^25–27^ and chondrocytes ^28^. The conventional application of SAP-based hydrogels in cell culture involves dispensing a mixture of cells and SAP solution onto the bottom of a culture well, followed by the addition of medium on top. This induces β-sheet formation, leading to in situ gelation and encapsulation of the cells. However, due to the inherently weak mechanical properties of SAPs, this method often results in inconsistent gel formation and the detachment of the hydrogel in sheet-like fragments, particularly during medium exchanges, thereby introducing heterogeneity and variability that significantly limit the reliability of SAPs in cell culture and screening applications.

Here, we present a novel protocol that employs SAPs to generate defined and scalable hiPSC-derived spheroidal 3D models, which we refer to as “*organobodies*” (OBs). Following extensive characterisation, we demonstrate that PSC-derived hepatocyte-like cells (HLCs) within the OBs exhibit a relatively mature hepatic phenotype, are metabolically active, and can detect the hepatotoxicity of a selected panel of chemical compounds to an extent comparable to that observed in 3D primary human hepatocytes (PHHs).

## Materials and Methods

### Cell Line Generation and Maintenance

We used three sources of human induced pluripotent stem cells (hiPSCs) in this study. The Sigma 0028 line was purchased form Sigma-Aldrich and genetically engineered to inducibly overexpress 3 transcription factors (TFs), namely *PROX1*, *FOXA3,* and *HNF1A* (the line is referred to as HC3X) as described earlier ^29^ to generate HLCs. HC3X cells were expanded feeder free on Matrigel (BD Biosciences) coated plates in Essential 8 Flex (Thermo Fisher Scientific). All studies using HC3X cells were approved by the Human Ethics Committee at the University Hospital, Gasthuisberg, KU Leuven, Belgium (S50354).

The lines UTA.10211.EURCCs (referred to as 10211) and UTA.10802.EURCCs (referred to as 10802) were generated in-house directly from fibroblasts derived from two individuals. Pluripotency was induced using the Sendai virus-based reprogramming kit (CytoTune; Life Technologies), which includes the transcription factors OCT4, SOX2, KLF4, and c-MYC, following protocols established by Takahashi and Yamanaka ^30^ and Ohnuki, Takahashi, and Yamanaka ^31^. The 10211 and 10802 lines were maintained on mitotically inactivated mouse embryonic fibroblasts (MEFs; #ASF-1223, Applied StemCell) as described earlier^32^. The study was approved by the ethical committee of Pirkanmaa Hospital District (R12123) and written consent was obtained from all fibroblast donors.

### hiPSC Hepatic Differentiation

hiPSCs HC3X were differentiated towards HLCs as described earlier ^29,33^. Briefly, PSCs were detached using StemPro^TM^ Accutase® Reagent (#A1110501, Thermo Fisher Scientific) and plated at 2 × 10^5^ cells/ml in mTeSR medium (#85850, Stem Cell Technologies) supplemented with 10 ul/ml RevitaCell (#A2644501, Thermo Fisher Scientific). The next day, differentiation was started in liver differentiation medium (LDM) until day 12. From day 12 to 40, a 3× concentrate of non-essential amino acids (AAs) (#11140035, Thermo Fisher Scientific), and from day 14 onward, 20 g/L glycine (#G8898, Sigma) was added to the LDM medium as described in Boon et al. ^29^. 5 µ/ml doxycycline was added from day 4 onward to induce the overexpression of *PROX1*, *FOXA3,* and *HNF1A*. Growth factors were added as follow: 50 ng/ml Activin A day 0 to day 4, 50 ng/ml Wnt-3a (day 0 to day 2), 50 ng/ml BMP4 (day 4 to day 8), 50 ng/ml aFGF (day 8 to day 12) and 20 ng/ml HGF (day 12 onward). All growth factors were purchased from Peprotech. DMSO was added to the medium with concentration of 0.6 % (between day 0 and 12) or 2% (between day 12 and 14). The differentiation was continued until day 20, 37-40, and 60.

Differentiation of hiPSCs 10211 and 10802 was adapted from the previously described protocol ^34^. Briefly, hiPSCs were detached by using Versene, and resuspended in DE induction medium consisting of RPMI + Glutamax supplemented with 1× B27, 100 ng/ml Activin A, 50 ng/ml Wnt3, and 10 μM ROCK inhibitor and seeded at a density of 5–10 × 10^4^ cells/cm^2^. After 24 hours, the ROCK inhibitor was replaced by 0.5 μM sodium butyrate (NaB), which was maintained until day 3–5 of DE differentiation. Then medium was changed to KO-DMEM medium supplemented with 20% KnockOut™ Serum Replacement (KO-SR), 1 mM Glutamax, 1% non-essential amino acids (NEAA), 0.1% β-mercaptoethanol (β-ME), and 1% dimethyl sulfoxide (DMSO) for 6–7 days. The final maturation phase was initiated by switching Hepatocyte Basal Medium (HBM; cc-3199, Lonza) supplemented with SingleQuots™ (Lonza), 25 ng/ml hepatocyte growth factor (HGF; #PHG0254, Life Technologies), and 20 ng/ml oncostatin M (OSM; #295-OM, R&D Systems). The maturation stage was continued for up to 26 days, with medium changes every other day.

As a reference for OBs generated from HC3X, primary human hepatocytes (PHHs) from three cadaveric donors F108 (male, 47), F125 (male, 62), and F133 (male, 57) were obtained from the Ministry of Health-accredited tissue banks at Cliniques Universitaires St. Luc, Brussels, under Belgian "opt-out" organ donation legislation. PHHs were cryopreserved post-isolation and used immediately after thawing as positive controls for hepatocyte marker expression and function. Cryopreserved PHHs (Cat# HMCPIS, Lot. HU8210, Gibco®) were used 1 day post plating as a reference for OBs generated from 10211 and 10802.

3D PHHs composed of pooled primary human hepatocytes provided by InSphero were maintained in Akura 96^TM^ spheroid microplates in 3D Insight^TM^ Human Liver Maintenance Medium-Tox (#CS-07-001-01, InSphero) for 4 days until cell toxicity studies.

HepG2 cells (ATCC) were cultured in RPMI + GlutaMAX™ medium (#61870, Gibco) supplemented with 10% FBS and used between passages 5–16, 2–5 days post-passaging.

### Generation of Organobodies

Hepatic progenitors derived from the HC3X line were detached on day 8 of differentiation using TrypLE Express (#A2644501, Thermo Fisher Scientific) for 8 min at 37 °C. Single cells were resuspended in LDM supplemented with 10% FBS, counted, and centrifuged at 300 × g for 4 min. The cell pellet was then resuspended in 10% sucrose prepared in Milli-Q water and centrifuged again. The pellet was then resuspended in 10% sucrose to achieve a final cell density of 2–6 × 10⁴ cells/µl. Cell suspension was kept on ice until further use.

RADA16 (commercially available as PuraMatrix, #354250, Corning) was vortexed vigorously for 2 min to reduce viscosity, then mixed with 10–20% collagen I (#A1064401, Thermo Fisher Scientific) before being mixed with the cell suspension at a 1:1 ratio in a 1.5 ml Eppendorf tube by gentle pipetting and stirring, avoiding bubble formation. The tube was kept on ice until use. A 6, or 96-well plate was pre-filled with day 8 differentiation medium supplemented with RevitaCell. Using an automatic pipette (0.5-20 ul), 1–7 µl of the SAP-cell mixture was carefully dispensed onto the surface of the medium to form discrete cell-gel droplets either in bulk (several droplets in 1 well of 6 wp maintained on a shaker) or individual (1 droplet per well of 96 wp) format. After 30 min, the medium was replaced with fresh medium containing RevitaCell or Rock inhibitor for 24 hrs before refreshing the medium. Medium change was continued according to the protocol for 2D culture.

Hepatic progenitor of cell lines 10211 and 10802 at day 8-10 of differentiation were dissociated using Gentle Cell Dissociation Buffer (#07174, STEMCELL Technologies) for 25 min or TrypLE™ Express (#12563011, Gibco™) for 5–7 min and a final cell pellet was generated in 10% sucrose as mentioned above with a density of 1.5–3 × 10^4^ cells/µL and mixed in a 1:1 ratio with PuraMatrix™ containing 20% bovine collagen I. Droplets (1–5 µL) of the cell–hydrogel mixture were manually dispensed using a 10 µL pipette directly into the HCM maturation medium (see Figure S1). Cultures were maintained under static conditions in HCM maturation medium for up to 26 days and medium was refreshed every other day.

The same procedure was applied for HepG2 cells except that they were detached using 0.25% trypsin.

### Generation of Conventional 3D Spheroids

Hepatic progenitors derived from the HC3X line were detached and centrifuged as described above. Cells were resuspended in day 8 medium supplemented with 10% Matrigel and seeded with the density of 2 × 10⁴ cells/well in ultra-low attachment U bottom plates (#650970, Greiner). Medium was supplemented by RevitaCell for the first 24 hrs. Subsequent medium change and differentiation steps were carried our in the same manner as for 2D HLCs and 3D hepatic OBs.

### RNA Extraction and RT-qPCR

For OBs and conventional spheroids, 2-3 individual samples were pooled in one tube per condition. For 2D HLCs or HepG2s, a well of 24 well plate was used per condition. RNA was extracted using Qiazol reagent (#79306, Invitrogen) according to the manufacturer’s instructions and quality checked by nanodrop. Up to 500 ng of RNA was reverse transcribed into cDNA using the SuperScript™ III First-Strand Synthesis Kit (#11752050, Invitrogen). Gene expression was analysed using the Platinum™ SYBR Green qPCR SuperMix-UDG Kit (#11733038, Invitrogen) on a ViiA™ 7 Real-Time PCR System (Thermo Fisher Scientific). Primer sequences are listed in Supplementary Table 1. Ribosomal protein L19 (RPL19) served as the reference gene for normalization.

RNA extraction and qPCR were performed on cell lines 10211 and 10802 according to the method described earlier ^34^. Here, *GAPDH* was used as a reference gene for normalization.

### Live/Dead Assay

OBs were washed twice with PBS and treated with a mixture of 0.2 μM fluorescent calcein-AM staining live cells (green color), and 1 μM ethidium homodimer-1 staining dead cells (red color) from a purchased kit (#L3224, Invitrogen). After 30 min of incubation, OBs were washed twice with PBS and imaged by an Evos FL cell imaging microscope.

### Histology

OBs were washed in PBS and fixed for 30 min at room temperature (RT) in 4% (w/v) paraformaldehyde (PFA; Sigma-Aldrich), washed three times with PBS, and stored in PBS at 4 °C until paraffin embedding. Sections (5 µm) were prepared using a microtome (Microm HM 360, Marshall Scientific).

For Hematoxylin and Eosin (H&E), paraffin was removed with xylene, and sections were rehydrated through a graded ethanol series (100–70%, v/v). H&E staining was performed by sequential incubation in Harris Hematoxylin, acid alcohol, bluing reagent, and Eosin-Y. Stained sections were dehydrated, cleared in xylene, and mounted using DPX mountant (#06522, Sigma-Aldrich).

### Immunofluorescence Staining

#### Sections

Following deparaffinization and rehydration, antigen retrieval was performed by incubating sections in antigen retrieval solution (#S1699, Dako) at 98 °C for 20 min. Samples were permeabilized with 0.1% (v/v) Triton X-100 (#T8787, Sigma-Aldrich) in PBS for 20 min, then blocked with 5% (v/v) donkey serum (Dako) for 40 min, and subsequently incubated with primary antibodies diluted in Dako Antibody Diluent (DAD, #S202230-2, Dako) overnight at 4 °C. After washing with 0.1% Triton X-100, sections were incubated with secondary antibodies (1:500) for 45-60 min at RT. Finally, samples were washed and mounted with Prolong Gold antifade reagent with DAPI (#P36931, Life Technologies). Images were acquired using a Zeiss Axioimager microscope and processed in ImageJ.

#### Whole mount

Fixed OBs were transferred to a 48-well plate using a wide-bore pipette tip. Samples were blocked and permeabilized in 5% donkey serum, 0.5% PBST, and 2% DMSO for 5 hours at RT on an orbital shaker. Blocking buffer was then replaced with 250 µl of primary antibody diluted in DAD, and samples were incubated for two nights at 4 °C on an orbital shaker. OBs were then washed four times with 0.1% PBST for 10 min each at RT on an orbital shaker and then were incubated overnight at 4 °C in the dark with 250 µl of fluorescently labelled secondary antibody diluted in DAD. After four additional 10 min washes in 0.1% PBST, nuclei were stained with DAPI (1:2000 in PBS) for 1 hour at 4 °C in the dark. Samples were washed twice with PBS, transferred to a silicone mold, and optically cleared using RapiClear (#RC14700, SunJin Lab). OBs were then mounted with a coverslip and imaged using a Nikon C2 confocal microscope.

All primary and secondary antibodies used in this study are listed in Supplementary Table 2.

### Functional Assessments

Albumin secretion was measured using the Human Albumin DuoSet ELISA Kit, and alpha-1 antitrypsin (A1AT) levels were measured using the Human A1AT DuoSet ELISA Kit (both from R&D Systems), following the manufacturer’s instructions. Values were normalised to cell number, as determined using a NucleoCounter® NC-200™ Automated Cell Counter (ChemoMetec).

### Drug Toxicity Measurements

Hepatic OBs and 2D HLCs (day 40) generated from the HC3X line, 3D PHH microtissues (day 4), and HepG2 cells (day 5), all cultured in a 96-well plate format were exposed to seven different concentrations of various compounds with known hepatotoxicity profiles. Treatments were applied for 3, 7, or 14 days. For long-term cultures, doses were re-administered on days 3, 7, and 10. Each concentration was tested in 3–6 technical replicates. Cell viability was assessed by quantifying ATP levels using the CellTiter-Glo® 3D Cell Viability Assay (#G9682, Promega) and luminescence was measured using a Promega GloMax® Explorer plate reader. IC₅₀ values were then calculated by fitting dose-response curves using nonlinear regression analysis in GraphPad Prism.

### Drug Biotransformation

Phase I biotransformation was assessed for OBs generated from the HC3X line and 3D PHH microtissues by incubating cells for 4 and 24 hours with a cocktail of probe substrates selective for the major cytochrome P450 (CYP) isoforms at the following final concentrations: 10 µM phenacetin (CYP1A2), 10 µM diclofenac (CYP2C9), 10 µM bufuralol (CYP2D6), 50 µM chlorzoxazone (CYP2E1), 5 µM midazolam (CYP3A4), and 10 µM bupropion (CYP2B6). Phase I metabolite formation in culture supernatants was quantified by high-performance liquid chromatography-tandem mass spectrometry (HPLC-MS/MS), as described ^35^. Values were normalised to protein content, measured by Pierce™ BCA Protein Assay Kit (#23227, Thermo Fisher Scientific).

### Bulk RNA Sequencing Analysis

RNA was extracted using QIazol reagent (Invitrogen) according to the manufacturer’s instructions and quality assessed by nanodrop. RNA integrity was assessed using Agilent 5400. Samples with RNA integrity number (RIN) of ≥ 5 were used for library preparation. Messenger RNA was purified from total RNA through affinity capture using poly-T oligo-conjugated magnetic beads. After fragmentation, libraries were prepared using the Novogene NGS Stranded RNA Library Prep Set (PT044) and checked for size distribution with the Agilent 5400 Fragment Analyzer. Quantified libraries were pooled and sequenced on Illumina NovaSeq X Plus platforms to generate 150 bp paired-end reads, aiming for a sequencing depth of approximately 30 million reads per sample.

Sequence reads were trimmed using BBDuk (ver. 39.08, RRID:SCR_016969), then mapped onto the Human GRCh38.p14 reference genome ^36^ available on ENSEMBL using the STAR aligner (ver. 2.7.11b) ^37^. Unique gene hit counts were calculated by using feature counts from the Subread package (ver. 2.0.7) ^38^; a comparison of gene expression between the groups of samples was performed using DESeq2 (ver. 1.44.0) ^39^ package for the R programming language. ComBat from the sva R package (ver 3.57.0) was used to adjust normalized counts from literature data for batch effects. Data was analyzed/plotted with the seaborn (ver. 0.13.2) ^40^, scikit-learn (ver. 1.7.1), numpy (ver. 1.26.4) ^41^, pandas (ver. 2.2.0) ^42^ libraries for the Python programming language. Functional enrichment analysis was performed by querying the STRING database ^43^ with reString (ver. 0.1.21) ^44^.

Principal component analysis (PCA) was performed using VST-adjusted, batch-corrected, and normalized count data. Principal coordinates were computed based on the top 2,000 most highly expressed genes. Batch correction was performed using the ComBat function from the sva R package (version 3.57.0) to adjust normalized counts and account for batch effects.

KEGG pathway enrichment analysis between OBs and 2D HLCs was conducted using the list of all differentially expressed genes with a p-value less than 0.05.

KEGG pathway enrichment analysis between OBs and organoids was performed using the subset of differentially expressed genes that met the following criteria: adjusted p-value < 0.05 and absolute log₂ fold change ≥ 2.5. To ensure interpretability, the analysis was limited to the top 2,000 differentially expressed genes that satisfied these thresholds.

### Statistical Analysis

For RT-qPCR, statistical analysis was performed using One-way ANOVA using Brown-Forsythe and Welch tests to account for unequal variances. For Drug biotransformation and ELISA, unpaired t-test with Welch’s correction was used. P < 0.05 was considered statistically significant.

## Results

### Droplets of SAP Supports the Growth and Differentiation of Hepatic Cells

In this study, we describe a novel and simple technique to generate spheroidal hydrogels from SAP droplets. We further demonstrate that hiPSC-derived hepatic progenitor cells can be successfully cultured within these droplets through a straightforward sequence of steps to form spheroidal cultures (Figures 1A and S1A). We initially tested this approach with hepatic progenitors generated from two hiPSC lines 10211 and 10802 following the protocol published by Kiamehr et al ^45^ and then extensively applied the technique to generate 3D cultures from a genetically engineered line (HC3X) in which the hepatic differentiation is directed via the doxycycline-inducible expression of 3 hepatic transcription factors, *PROX1*, *FOXA3,* and *HNF1A* in an amino acid-rich medium ^29^. We showed that an even distribution of hepatic progenitors encapsulated in SAP droplets was achieved and a uniform growth was observed throughout the differentiation by using bright field microscopy (Figures 1B and S1C). Using a live/dead assay, we confirmed that the hepatic progenitors remained viable in SAP droplets, with minimal cell death observed (Figure S1D), indicating a cell-friendly encapsulation process.

**Figure 1.**
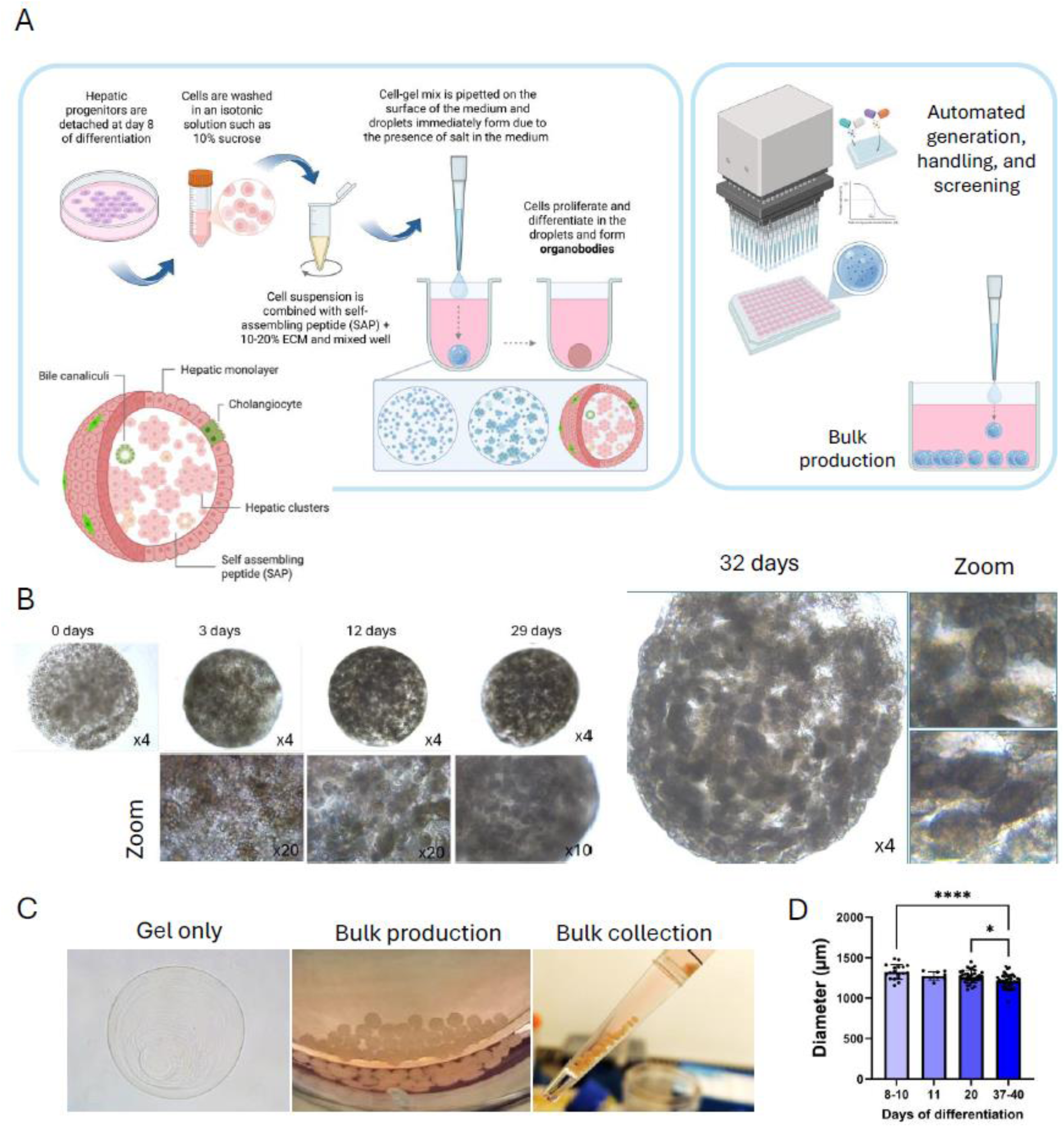
Generation and morphological observation of *organobodies* (OBs). **A)** Schematic overview of the process for creating OBs using self-assembling peptides from hiPSC-derived hepatic progenitors trough SAP droplet generation. The right panel illustrates the scalability of this approach, showing its compatibility with both automation and bulk production. *Image created with BioRender.com*. **B)** Brightfield images of hepatic 3D SAP droplets 0, 3, 12, 29, and 32 days after encapsulation of day 8 hepatic progenitors differentiated from HC3X line. **C)** Left: Brightfield image of an SAP droplet without cells. Middle: Bulk generation of OBs in a single well. Right: bulk transfer of maturated hepatic OBs using a 10 mL pipette. **D)** Quantification of OB dimensions over time, starting from day 8 of hepatic differentiation (0 days post-encapsulation) to day 40 (32 days post-encapsulation), with droplets prepared at an initial volume of 3 µL. Measurements were based on diameters from 110 brightfield images across three biological batches. Data are shown as mean ± SD. Statistical analysis was performed using unpaired two-tailed t-tests. P < 0.05 was considered statistically significant. *p < 0.05; ****p < 0.0001.

The size of the spheroidal cultures could be easily adjusted by modifying the initial pipetting volume of the SAP–cell mix droplets (Figures S1B and S3B). The resulting spheroidal structures were highly uniform, with a size variation of less than 7% (Figures 1C and 1D). Notably, the spheroidal cultures decreased in size by 8.8% during hepatic differentiation compared to their initial dimensions, suggesting biodegradation of the SAP nanofibers.

The culture density could be adjusted by varying the initial cell concentration. We started with a low density of 10,000 cells/µl (Figure S2) and increased it up to 30,000 cells/µl (Figure 2), which resulted in a corresponding increase in both the number and size of cell clusters within the droplets, contributing to a more tissue-like architecture in the 3D culture (Figure 2A).

**Figure 2.**
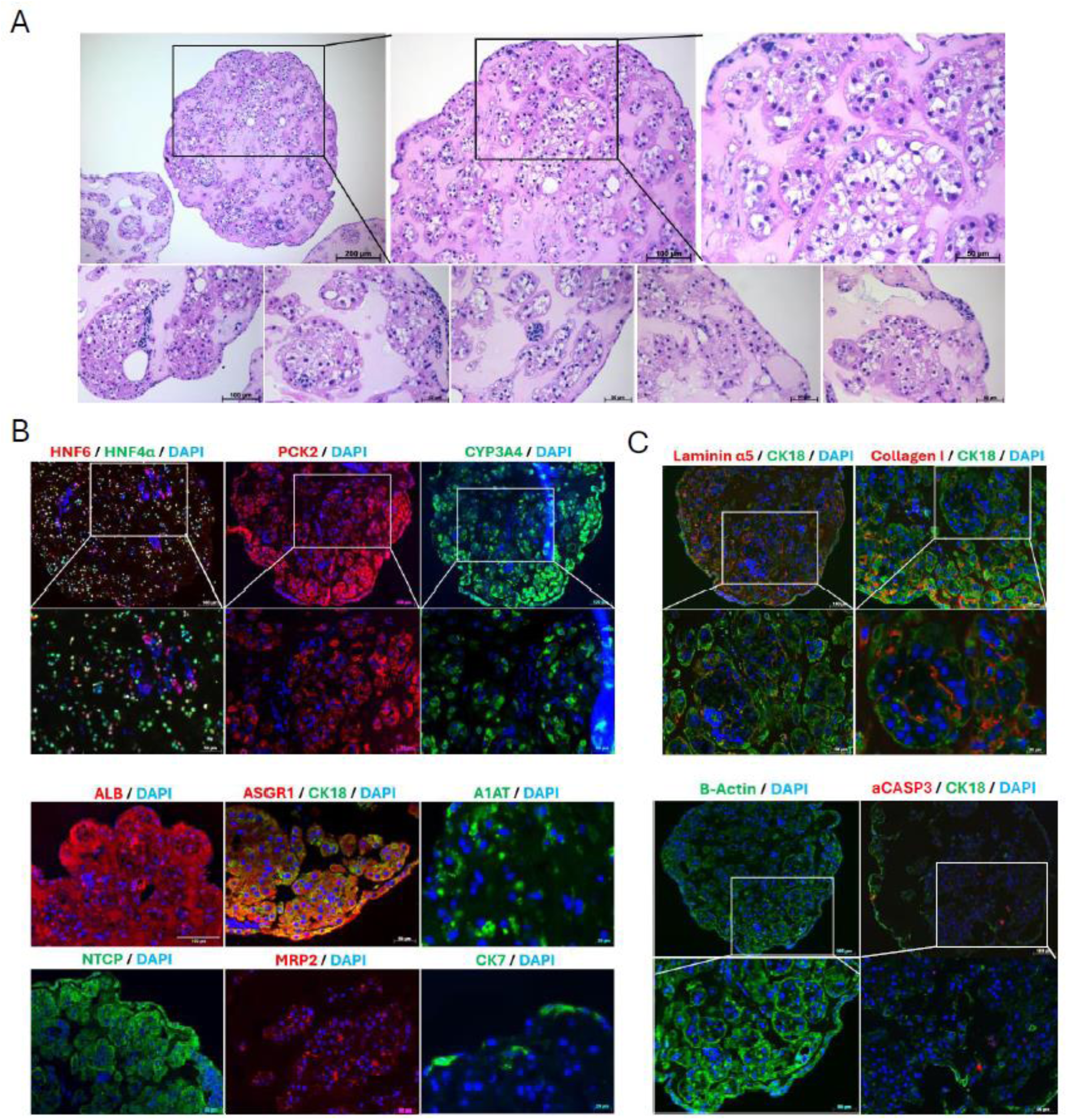
Histological and immunohistochemical characterization of hepatic OBs. **A)** Hematoxylin and eosin (H&E) staining of hepatic OBs 13 days after encapsulation of day 8 hepatic progenitors derived from the HC3X line, showing morphology, cellular distribution, and structural organization of HLCs. **B)** Immunohistochemical staining of OB sections 13 days post-encapsulation, demonstrating expression of key hepatocyte-specific markers: HNF4α, HNF6, CK18, PCK2, CYP3A4, ALB, ASGR1, A1AT, NTCP, and MRP2. The cholangiocyte marker CK7 was also detected. **C)** Immunostaining for ECM components laminin α5 and collagen I, and structural protein β-actin, revealed organized HLC clustering within the SAP hydrogel, indicative of matrix remodelling and ECM deposition. Staining for activated caspase-3 (aCASP3) confirmed minimal apoptosis within OBs. Nuclei were counterstained with DAPI. All images are representative of at least three independent experiments.

We also investigated the optimal timing for transition from 2D to 3D culture by harvesting differentiating HC3X progeny detached at various stages of hepatic differentiation. Our results showed that cells detached on day 8 (corresponding to the early hepatic progenitor stage) yielded the best outcomes in terms of final cell density, cluster size, and hepatic phenotype (Figure S3A).

Following encapsulation, hepatic progenitors proliferated and underwent morphogenesis, remodelling their environment by connecting with one another to form a tissue-like architecture. During the maturation stage, a uniform monolayer of hepatocytes with polygonal morphology emerged, covering the entire surface of the SAP droplets (Figures S1C and S2A) and surrounding clusters of HLCs formed within the droplets. From this point onward, we refer to these enclosed organelle-like structures as *organobodies* (OBs). H&E staining confirmed the formation of cellular clusters with typical hepatic morphology, along with a peripheral monolayer enclosing the culture system (Figures 2A and S2A).

Extensive immunocytochemistry and immunohistochemistry revealed that HLCs in OBs exhibit a mature phenotype, as confirmed by positive expression of key hepatocyte-specific markers, including ALB, HNF4α, HNF6, PCK1, PCK2, NTCP, CK18, ASGR1, and MRP2, indicating that the SAP droplet culture system effectively supports HLC maturation (Figures 2B, S2B–C). Notably, in prolonged cultures (5–6 weeks), HLCs began to form lumen-like structures with strong expression of CK18 and MRP2, suggesting advanced microenvironmental reorganisation and polarization of HLCs in these regions (Figures S2A and S2C). Additionally, a population of CK7-positive cells was observed, indicating the presence of cholangiocytes within the OBs. Interestingly, this cholangiocyte population was primarily located near the peripheral monolayer or within lumen-like areas inside the OBs (Figures 2B and S2D). β-actin expression showed structural orientation of HLC clusters connected within the OBs. We then assessed the deposition of two common hepatic ECM components, collagen I and laminin α5, and found that they were localized adjacent to the hepatic clusters, indicating matrix remodelling and structural reorganisation of the ECM by hepatic cells (Figure 2C). Despite the relatively large size of the OBs (∼1.3 mm), no necrotic core was observed, as confirmed by aCASP3 staining.

RT-qPCR analysis performed on days 20 and 37–40 (prolonged cultures) demonstrated that HLCs in OBs exhibited enhanced maturation compared to those maintained in 2D. As early as day 20, we observed significantly higher expression of key hepatic genes, including *PEPCK1*, *HNF4* α, *AFP*, *G6PC*, *CYP3A4*, *CYP2C9*, *CYP2C19*, and *CYP2D6* in OBs compared to the 2D HLCs. Further maturation was evident in prolonged cultures, where hepatic OBs expressed *ALB*, *PCK1*, *HNF4α*, *CYP3A4*, and *CYP3A5* at levels comparable to freshly isolated primary human hepatocytes (PHHs). Nevertheless, similar to 2D HLCs, *AFP* expression remained elevated in OBs compared to PHHs, suggesting that not all HLCs in OBs had yet reached a fully mature hepatic profile (Figure 3A).

**Figure 3.**
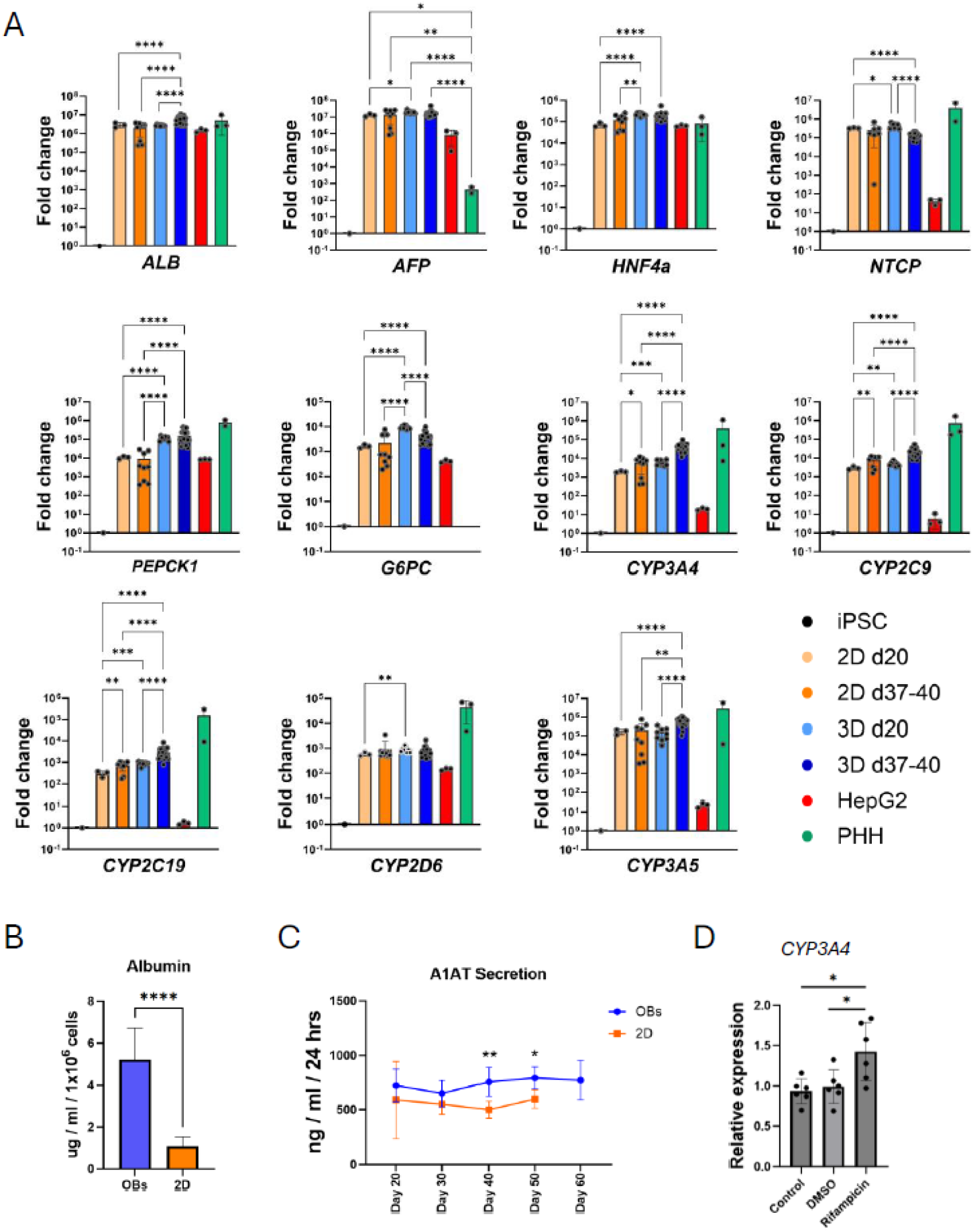
Gene expression profiling and functional characterisation of hepatic OBs. **A)** Comparative RT-qPCR analysis between 2D HLC cultures (*N* = 1-3 independent differentiations, *n* = 3-9 technical replicates) and 3D hepatic OBs (*N* = 3-7 independent differentiation, *n* = 9-21 technical replicates) generated from HC3X line for 11 key markers at early (day 20) and late (days 37 – 40) stages of differentiation in comparison with 2D HepG2 cells and freshly isolated PHHs (from 2-3 individual batch). *RPL19* was used as housekeeping gene to normalise gene expression. Fold change has been calculated against the expression of the same genes in undifferentiated HC3X hiPSC. To avoid overcrowded graphs, hiPSC and HepG2 cells have been excluded from statistical analysis. **B)** Albumin secretion levels in OBs versus 2D HLCs at day 40 of differentiation, measured by ELISA (N = 3; n = 9–12). **C)** A1AT secretion in the culture medium by hepatic OBs and 2D HLC cultures at days 20, 30, 40, 50, and 60 of differentiation measured by ELISA. The secretion level is significantly higher for OB cultures at days 40 and 50. (*N* = 3, *n* = 9-12). **D)** qPCR analysis showing significant increased expression of *CYP3A4* in 3D hepatic OBs following induction with 10 μM Rifampicin compared to vehicle (DMSO) and untreated control (*N* = 2, *n* = 6). Statistical significance was determined at *P* < 0.05. *\*P < 0.05; **P < 0.01; ***P < 0.001; ****P < 0.0001*. Data are presented as mean ± SD.

Our comparative study with classical spheroids showed that OBs expressed higher levels of several hepatocyte-specific markers, including *CYP3A4*, *CYP1A2*, *HNF4α*, and *PEPCK* (Figure S6).

### OBs Demonstrate High Levels of Functionality

From this point onward, our functional analysis focused exclusively on HLCs derived from the HC3X cell line. Compared with parallel 2D HLC cultures, OBs secreted significantly higher amounts of albumin and A1AT into the culture medium, with differences becoming significance on days 40 and 50 (Figures 3B–C), indicating superior maturation of HLCs in OBs. Furthermore, stable A1AT secretion from day 20 to day 60 demonstrated the long-term stability of the OB culture system, consistent with RT-qPCR data showing sustained hepatic gene expression over the same period (Figure S3D).

To assess functional responsiveness, hepatic OBs were treated with 10 µM Rifampicin for 48 hours, resulting in a significant induction of *CYP3A4* expression, confirming the maturity and metabolic competence of the HLCs within the OBs (Figure 3D).

### OBs Exhibit Higher Maturity than 2D HLCs

Through comprehensive analysis, we explored the gene expression profile of OBs (N=5). HepG2 cells (N=3), and PHHs (N=3) as well as iPSC-derived organoids as reported in Shimizu et al. ^46^ (GEO accession: GSE248110 and GSE247960), and PHHs reported in Boon et al. ^29^ (GEO accession: GSE140520), and Marsee et al. ^47^. Principal-component analysis (PCA) of the top 2000 features revealed distinct gene expression profiles for OBs, organoids, and PHHs, with each forming separate clusters (Figure 4A). PC1 primarily separated PHHs from OBs, organoids, and HepG2 cells, while PC2 positioned OBs furthest from HepG2 cells, with organoids exhibiting an intermediate phenotype between OBs and HepG2 cells. Heatmap analysis of differentially expressed genes identified more than 90 genes that were significantly different between OBs and 2D HLCs at day 37–40 of differentiation (Figure 4B). KEGG pathway analysis of 1,353 differentially expressed genes (p < 0.05) revealed significant enrichment of several biological processes, including multiple metabolic pathways such as drug metabolism–cytochrome P450 (Figure 4C). Heatmaps visualization of genes within the top enriched pathways demonstrated upregulation of CYPs (*CYP3A4*/*5*, *CYP2C9*, *CYP1A2*) and phase II enzymes (*UGT2B11*, *UGT2B7*) in OBs compared with 2D HLCs, highlighting enhanced metabolic competence and maturation of OBs (Figure 4D). Additionally, *ADH1B*, a key gene involved in alcohol metabolism in the adult liver, was expressed at higher levels in OBs than in 2D HLCs, further supporting the superior maturity and predictive potential of OBs for toxicity screening. Genes associated with PPAR signalling, including *CD36*, *HMGCS2*, and *PCK1*, were also more highly expressed in OBs, indicating enhanced lipid uptake, ketogenesis, and gluconeogenesis in OBs, features strongly associated with metabolic maturity. Moreover, *RXRA*, an important transcription factor in hepatocyte differentiation and maturation ^48^ was expressed higher in OBs. KEGG pathway analysis also revealed enrichment of fatty acid (FA) metabolism in OBs compared with 2D HLCs.

**Figure 4.**
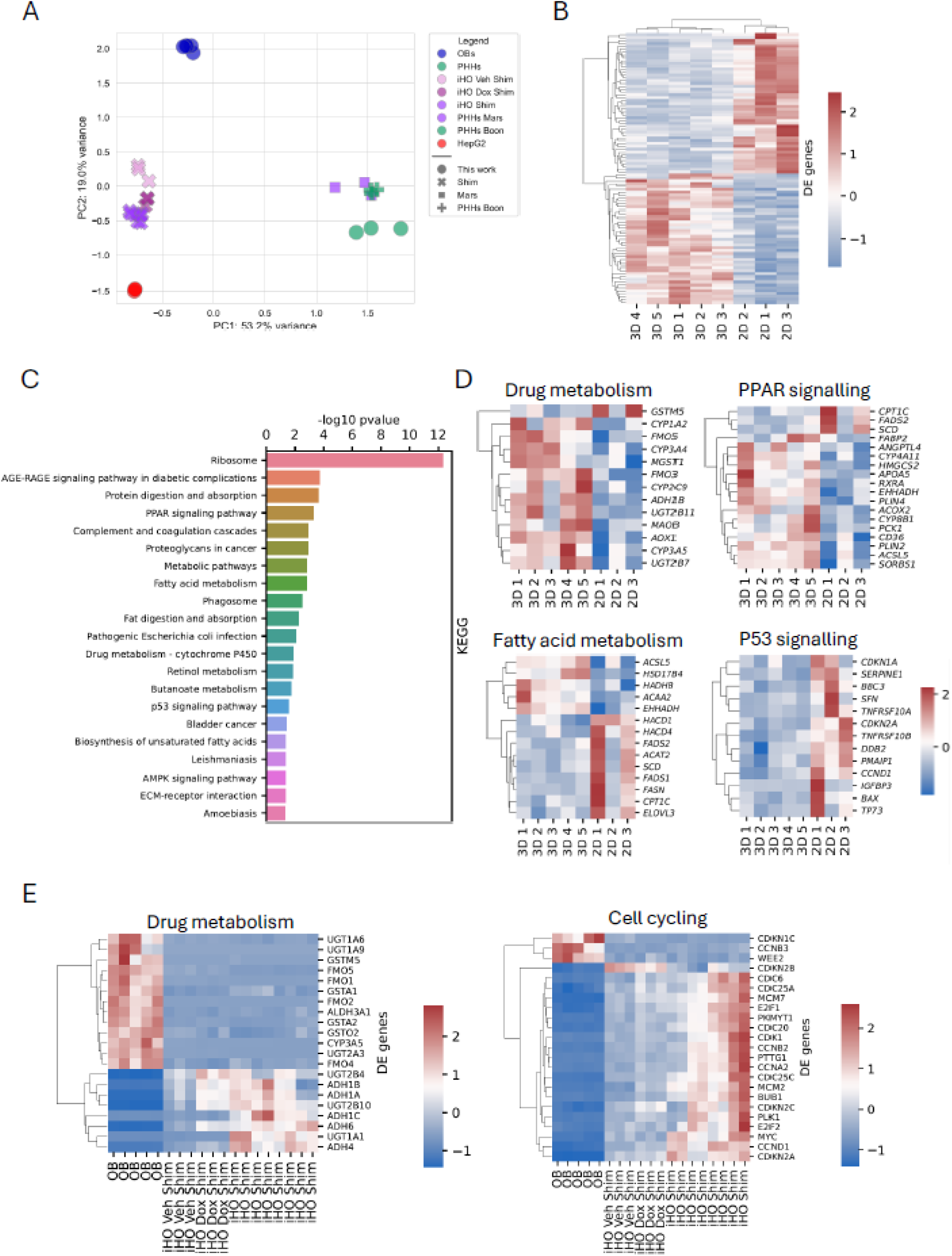
Bulk RNA-sequencing analysis of hepatic OBs. **A)** Principal-component analysis (PCA) comparing global expression profiles of OBs (N=5), HepG2 cells (N=3), and freshly isolated PHHs (N=3) as well as iPSC-derived organoids as reported in Shimizu et al. ^46^ (GEO accession: GSE248110 and GSE247960), and PHHs reported in Boon et al. ^29^ (GEO accession: GSE140520), and Marsee et al. ^47^ (GEO accession: GSE298576). **B)** Heatmap showing the differentially expressed (DE) genes between OBs (3D) and 2D HLCs. Values are represented as log2 fold changes, with red indicating genes upregulated and blue indicating genes downregulated. Each row represents a gene, and each column represents a sample from individual experiments. **C)** Bar plot of the enriched KEGG pathways based on all DE genes between OBs and 2D HLCs with a p-value less than 0.05 (1353 genes). Bar length corresponds to -log₁₀ p-values. **D)** Heatmaps of bulk RNA-seq data representing the z-score values across OBs (3D) and 2D HLCs showing genes associated with drug metabolism, PPAR and P53 signalling pathways as well as fatty acid metabolism. **E)** Heatmaps of bulk RNA-seq data representing the z-score values across OBs (3D) and iPSC-derived hepatic organoids (iHO) showing genes associated with drug metabolism and cell cycling.

Notably, genes involved in FA β-oxidation (*HADHB*, *ACAA2*, *EHHADH*, *HSD17B4*) were upregulated in OBs, whereas genes associated with lipogenesis and FA elongation/desaturation (*FASN*, *SCD*, *FADS1/2*, *ELOVL3*) were downregulated. These expression changes indicate a metabolic shift from lipogenesis toward FA catabolism in OBs. In contrast, genes linked to the P53 signalling pathway, including *BBC3*, *DDB2*, *BAX*, *TNRSF10A* and *TNRSF10B*, were expressed at lower levels in OBs, suggesting that cells in OBs experience less stress compared with 2D cultures.

Comparison of OBs with iPSC-derived hepatic organoids (iHOs) from a publicly available dataset revealed comparable drug-metabolizing capacities between the two groups of samples (Figure 4E). Notably, the cell cycle pathway was among the top enriched pathways, with most cell cycle–related genes upregulated in iHOs, suggesting that these cells have retained a more proliferative state compared to OBs.

### Hepatic OBs Accurately Predict Drug-Induced Liver Injury and Exhibit Functional CYP-Mediated Drug Metabolism

We also investigated whether hepatic OBs could be used to detect drug-induced liver injury (DILI). OBs at day 40 were exposed to 13 different compounds representing low (Buspirone, Entacapone, Rosiglitazone), moderate (Clozapine, Acetaminophen, Obeticholic acid), and high (Troglitazone, Tolcapone, Nefazodone, Trovafloxacin) DILI risk according to the FDA DILI severity ranking ^49^. Total cellular ATP content, as an indicator of relative cell viability, was measured after 3, 7, or 14 days of treatment. We compared the performance of OBs with 2D HepG2 cells, widely used in the pharmaceutical industry, 2D HLCs, and gold standard hepatic 3D microtissues from InSphero.

Next, we compared hepatic OBs, 2D HLCs, and 3D PHH microtissues for three compounds (Tolcapone, Entacapone, and Troglitazone) under 3- and 7-day dosing regimens. OBs outperformed 2D HLCs in the short-term (3-day) treatment, as the highest concentration (480 µM) of Tolcapone and Entacapone caused only 89% and 75% ATP reduction in 2D HLCs, respectively, without complete cell death. In contrast, 2D HLCs exhibited greater sensitivity to Troglitazone after 3 days compared to the other two models. Troglitazone is a lipophilic compound, and its higher cytotoxicity in 2D HLCs may be explained by their lipid-rich phenotype. As will be shown later in the results, bulk RNA-seq analysis indicates that 2D HLCs are more lipogenic than OBs, suggesting a higher intracellular lipid content. This lipid-rich environment likely facilitates the dissolution and accumulation of lipophilic compounds such as Troglitazone within cellular membranes and lipid droplets, leading to elevated intracellular drug levels and, consequently, greater cytotoxicity.

Alternatively, OBs and 3D PHH microtissues showed comparable performance in predicting the toxicity of Tolcapone, Entacapone, and Troglitazone (Figure 5B). After 7 days of dosing, all three models exhibited similar sensitivity to the tested compounds. Interestingly, we could also morphologically observe the response of OBs to toxicity: toxic compound concentrations affected the smooth peripheral monolayer, causing detachment of cells from the OB surface, which was easily visible under brightfield microscopy (Figure 5C).

**Figure 5.**
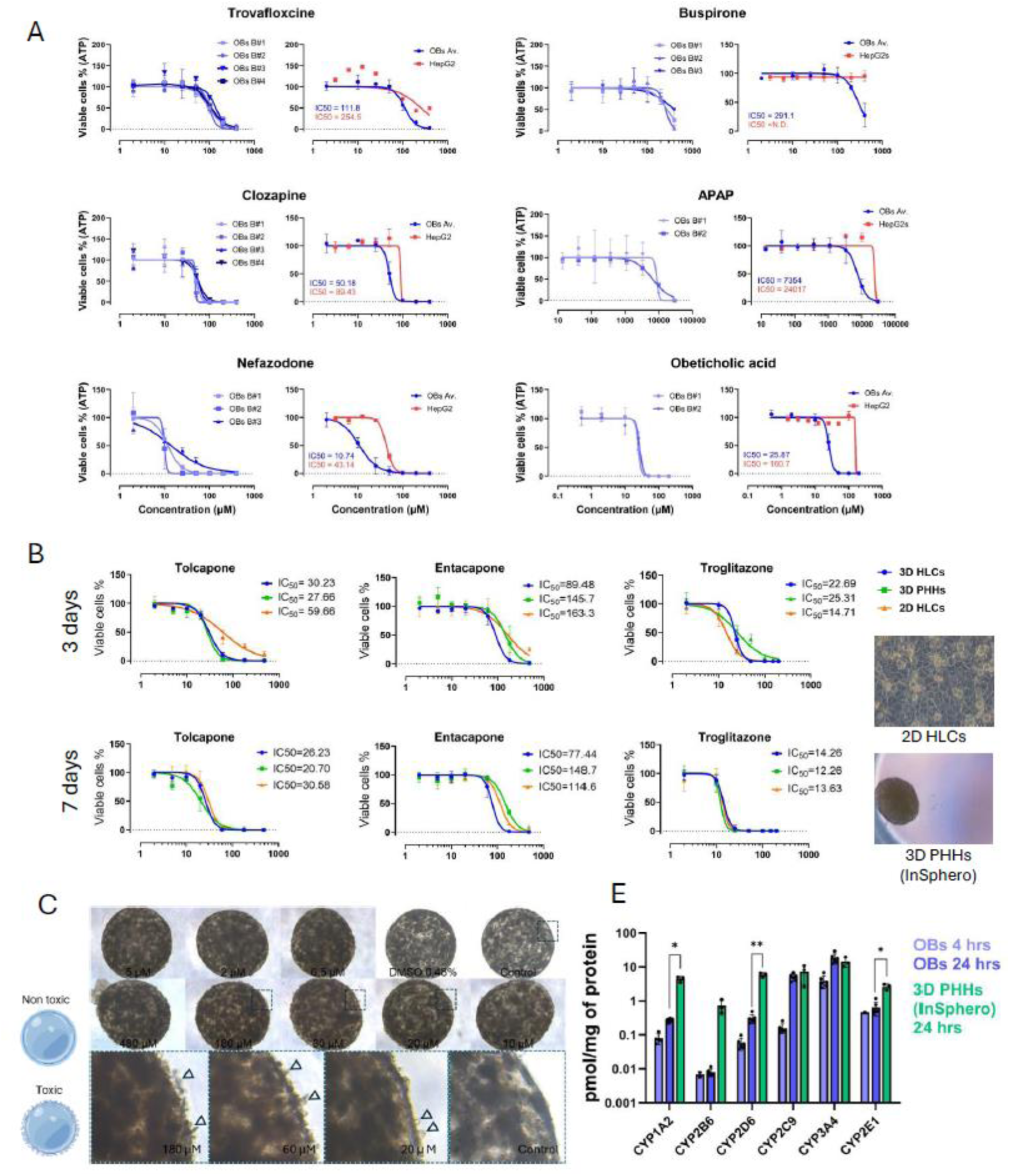
Comparative drug toxicity and metabolism profiling of hepatic OBs. **A)** Dose-response toxicity curves comparing 3D hepatic OBs and 2D HepG2 cells after 7 days of treatment with Trovafloxacin, Clozapine, Nefazodone, Buspirone, Acetaminophen (APAP), and Obeticholic acid. Left panels show raw data for individual biological replicates (N = 2–4, with 3–6 technical replicates per concentration). Right panels display averaged dose-response curves with corresponding IC₅₀ values, demonstrating superior sensitivity of OBs over HepG2 cells. **C)** Comparative dose-response curves for OBs (N = 3 biological repeats, n = 4 technical replicates), 2D HLCs (N = 1, n = 3), and 3D PHH microtissues (N = 1, n = 3) following 3-day (top row) and 7-day (bottom row) exposure to Tolcapone, Entacapone, and Troglitazone. IC₅₀ values were calculated using nonlinear regression analysis (GraphPad Prism). **C)** Brightfield images showing morphological changes in OBs treated with increasing concentrations of Tolcapone. Arrows indicate cell detachment from the spheroid surface, consistent with observed toxicity (panel B). **D)** Drug metabolism capacity of OBs (N = 3, n = 6) compared to 3D PHH microtissues, assessed after 4 and 24 hours of treatment with a cocktail of six reference substrates, each metabolized by a different CYP enzyme. Metabolite formation was quantified by HPLC-MS/MS. Statistical comparisons were performed using unpaired t-tests with Welch’s correction.

We then benchmarked the drug biotransformation capacity of OBs against gold standard 3D PHH microtissues by exposing the hepatic cultures to a cocktail of 6 compounds, each primarily metabolized by a specific CYP enzyme: acetaminophen (CYP1A2), hydroxybupropion (CYP2B6), hydroxybufuralol (CYP2D6), 4′-hydroxydiclofenac (CYP2C9), 1′-hydroxymidazolam (CYP3A4), and 6-hydroxychlorzoxazone (CYP2E1). CYP3A4- and CYP2C9-mediated drug biotransformation appeared comparable between OBs and 3D PHH microtissues, consistent with our RT-qPCR data where we compared OBs with three batches of fresh 2D PHHs. In contrast, the activities of other tested enzymes were lower in OBs than 3D PHH microtissues, which was statistically significant for CYP1A2, CYP2D6, and CYP2E1. This suggests that OBs still have limited biotransformation capacity for certain CYP pathways compared to PHHs and improving HLC maturation will be necessary to reach full drug metabolization potential.

## Discussion

Liver organoids hold great potential for drug hepatotoxicity screening and the evaluation of drug-induced liver injury (DILI) ^50^. However, developing robust and scalable models capable of preserving liver-specific functions over the long term remains technically challenging ^51^. In this study, we developed a novel technique based on a synthetic self-assembling peptide (SAP) hydrogel, for the generation of robust and size-controllable liver organoids. This simple, defined, and minimal approach enables encapsulation of cells in a spheroidal format within seconds, triggered by the natural salt content of the culture medium. Using this technique, we were able to generate and maintain functional and mature 3D hepatic cultures that were stable for several weeks in culture. Our SAP-based culture system avoids the need for undefined animal-derived matrices or time-consuming gelation steps, ensuring reproducibility and uniformity, and is amenable to scalable, automated, and high-throughput drug toxicity screening. Additionally, in the OB model, HLCs acquire their polarity, maturity, and extensive drug metabolizing activity, demonstrating a consistent and robust ability to predict the toxicity of several hepatotoxicants, comparable to “gold standard” 3D primary hepatocytes.

Reliance on undefined matrices such as Matrigel is a critical contributing factor to the variability and heterogeneity of 3D and organoid systems, and its undefined composition also severely limits Matrigel in clinical studies ^10,17,52–57^. Several defined hydrogels, such as nanofibrillar cellulose ^57^, gelatin ^58^, Polyisocyanopeptides (PIC) ^59^, or functionalised PEG ^60^ hydrogels have been developed for tuneable, large-scale production of classical cystic liver organoids from LGR5+ cells and iPSCs. However, unlike these defined hydrogels, organobodies (OBs) generated in SAP demonstrated high levels of CYP enzymes activity (particularly CYP3A4 and CYP2C9), albumin expression and secretion to similar levels as those in PHHs. Additionally, despite their value for creating patient-specific models, classical organoid culture systems are inherently challenging due to their limited proliferative capacity restricting long-term expansion ^61^. Furthermore, the lack of donor material, ethical constraints and biological heterogeneity could hamper further progress, particularly in scaled-up applications ^55^. Therefore, patient specific organoids developed from hiPSCs could be a better approach particularly for drug efficacy and toxicity screening ^9,46^. However, some challenges still remain such as achieving high and consistent levels of CYP expression, improving metabolic maturation, and reducing the variability between organoid batches addressing which would increase the relevance of organoids for predictive hepatotoxicity ^61–63^. Here, we generated OBs from three individual hiPSC lines in two independent laboratories by utilizing two different hepatic differentiation protocols. One of the cell lines that we previously extensively studied was a genetically engineered line (HC3X) equipped with 3 inducible transcription factors: *HNF1*, *FOXA3*, and *PROX1*. We recently demonstrated that inducible overexpression of these transcription factors during hepatic differentiation could boost hepatic differentiation and maturation in amino acid rich (AAGly) medium in 2D ^29^. By culturing HC3X HLCs in 3D SAP droplets, we could induce transcriptional and functional maturation of these HLCs even further. In fact, RNA-sequencing analysis revealed increased expression of drug-metabolizing enzymes and genes involved in the PPAR signalling pathway, indicating a more advanced maturation of hepatocytes in OBs compared with 2D HLCs. We also observed a coordinated upregulation of mitochondrial and peroxisomal β-oxidation genes alongside downregulation of de-novo lipogenesis and desaturation machinery suggesting that OBs promote a more oxidative, mature hepatocyte phenotype. This metabolic reprogramming is consistent with activation of PPARα-driven pathways, which underlines improved functional maturity in OBs. We also compared our findings with publicly available RNA sequencing data from an advanced organoid culture system enhanced through FXR signalling, recently published by Shimizu et al. ^46^. Unlike conventional organoids, which are typically characterized by cystic structures resembling bile ducts, the organoids (iHOs) generated by Shimizu et al. exhibited a grape-like morphology reminiscent of fetal hepatocytes. Although OBs and iHOs clustered separately in the PCA plot indicating their distinct gene profile, both demonstrated comparable drug-metabolizing capacities (Figure 4E). Notably, iHOs displayed higher expression of cell cycle-related genes, consistent with their maintained proliferative capacity, as indicated by positive Ki67 staining reported by the authors.

In this study, we utilised PuraMatrix, a commercial self-assembling peptide composed of RADA16-I sequence. The material properties of RADA16-I have been previously characterized ^21^ and its ability to support growth and differentiation of several mammalian cells has been already demonstrated in 3D ^27,64–66^. However, all the previous studies have followed the conventional method of dispensing a mixture of cells and SAP hydrogel onto the bottom of a culture well, followed by the addition of medium on top. While this approach is effective for hydrogels like Matrigel or collagen, it is problematic when SAP-based hydrogels are used due to hydrogel fragmentation causing sample loss and increased heterogeneity among technical replicates (Figure S7A). Here, we took a different approach, and instead directly pipetted the droplets of the SAP-cell mixture into the culture medium. As the droplet immerses in the medium at once, this leads to formation of stable spheroidal hydrogels remaining intact throughout long-term culture and repeated medium changes. Our approach not only tackled a challenge facing the applicability of SAP-based hydrogels but also resulted in a distinctive and unique culture system in a spheroidal shape, where a monolayer of polygonal and polarized hepatocytes covered the entire periphery of the spheroidal hydrogel creating a large lumen filled with SAP. Inside this lumen, hepatic clusters where formed that could merge to create larger clusters. This configuration distinguishes the OBs from both classical epithelial organoids generated from adult stem cell-derived LGR5+ cells ^53,59,60,67^ or 3D cellular aggregates generated from pluripotent stem cells such as iPSCs with a compact cellular organisation. We chose the term *Organobodies* (OBs) for this unique culture system as the SAP hydrogel provided a spheroidal “body” for generation of liver organoid-like structures.

In high-throughput assays such as toxicity and drug screening, achieving uniformity in the size, shape, density, and maturity of 3D cultures is critical for reducing inter- and intraplate/batch variation ^68^. While consistent shape and size ensure similar surface area-to-volume ratios and diffusion distances for drug penetration, uniform cell density assist with comparable baseline and metabolic activity levels, preventing variability in assay readouts such as cell viability. Additionally, uniform maturity across spheroids ensures consistent levels of drug metabolization capacity, which is particularly important for assessing compounds that require metabolic activation. One of the approaches in this direction has been generating organoids in microspheres or microbeads from alginate ^69–73^. However, despite its low cost, alginate is bio-inert, lacks cell-adhesive cues, and is naturally resistant to enzymatic degradation by mammalian cells. Using RADA16, we were able to produce spheroidal hydrogels with uniform size and shape resulting in repeatable batches of mature OBs. Even though the synthetic RADA16 hydrogel contains no biologically active domains for cell attachment, it has been shown that the high surface area-to-volume ratio created by peptide nanofibers provide spatial cues that promotes cell adhesion, proliferation, and differentiation ^26,74^. Despite this, we supplemented SAP with 10-20% collagen, which promoted cellular attachment and growth (data is not shown). Additionally, in the optimal culture environment, hepatocytes can secret their own ECM ^75,76^ and we could confirm the deposition of collagen I and laminin adjacent to the cellular clusters and beneath the monolayer periphery of the OBs critical in their morphogenesis, remodelling, and spheroid formation. Interestingly, we observed significantly higher ECM (collagen I and laminin α5) deposition in conventional spheroids made in ULA plates compared to OBs (Figure S6D and E). Collagen I is the main component of interstitial matrix and is found to increase with development of liver fibrosis ^77,78^. Laminin is mainly found in basement membrane and its correlation with liver fibrosis is less pronounced than collagen. However, studies have shown that in liver fibrosis, laminin deposition increases in space of Disse resulting in the formation of a perisinusoidal basement membrane leading to a process called capillarization ^79^. Additionally, laminin is providing support for proliferation and differentiation of hepatic progenitor cells in liver injury and fibrosis ^80^. These findings suggest that the SAP hydrogel used in our model might offer advantage in preventing or reducing fibrosis-like ECM remodelling. However, proving this requires further investigational studies. As the interaction of hepatocytes with other specific components of ECM may be beneficial in their maturation, it may also be of future interest to determine if further functionalizing the SAP with additional ECM/CAM peptides for an even better support of the hepatic cells.

Another approach for generating 3D liver constructs involves the formation of core-shell microcapsules, in which one or more cores are enclosed by an outer shell layer ^69,81,82^. These techniques often require sophisticated instruments such as coaxial flow or capillary microfluidics systems, involve lengthy or multistep sample handling, or typically rely on chemical or UV cross-linking and additional washing steps. The complexity of the 3D culture system is often directly associated with increased variability and heterogeneity of the final 3D models ^17^ and producing reproducible batches with similar mechanical properties, stability and permeability remains to be a challenge with these techniques. In our approach, SAP-cell mix can be directly pipetted or printed into the culture medium where the encapsulation happens in fraction of seconds simply triggered by the natural salt content in the medium, therefore removing the need for extra cross-linking or washing steps. In core-shell microcapsules, cells are mostly in the form of aggregates in the core and are hard to retrieve while the cells in OBs are evenly distributed and could be recovered by mechanical pipetting only and without enzymatic digestion of the SAP. In addition, it remains unclear whether the shell of core-shell microcapsules interferes with the penetration of large compounds into the core, where the cells reside. Admittedly, we did not directly investigate the diffusion rate of large compounds through OBs; however, we observed complete cell death at high concentrations of all tested drug compounds (Figures 5 and S5), suggesting that compound penetration was not limiting under our experimental conditions.

RT-qPCR analysis revealed enhanced maturity in OBs generated from all three hiPSC lines compared to parallel 2D HLCs. This maturation increased with prolonged culture and remained relatively stable until day 60 of culture. We also investigated whether the initial OB size influences hepatic gene expression by varying the volume of the SAP-cell mixture (1, 2, 3.5, and 7 µL). Although certain genes, such as *PEPCK1* and *CYP3A4*, showed differential expression across OBs of different sizes, we did not observe consistent or substantial changes in the overall hepatic gene expression profile across the OBs with various sizes (Figure S3D and E). Nevertheless, we found that the volume 3-5 µL results in optimal OBs in terms of handling and providing sufficient cells for downstream analysis. Applying smaller volumes bellow 1 µL was still feasible, however, more sophisticated instruments may be required for volumes less than 0.5 µL.

We observed that in 2D HLCs derived from lines 10211 and 10802, the expression of hepatic genes declined over prolonged culture, indicating de-differentiation of HLCs in this system. In contrast, OBs generated from the same lines maintained stable or even increased expression of hepatocyte-specific markers, such as *CYP3A4* (Figure S4). These findings highlight the importance of an optimal 3D microenvironment for sustaining prolonged and stable hepatic maturation in culture.

Importantly, we demonstrated that HLCs can maintain their hepatic phenotype for up to 60 days in culture without the need for passaging. Given that HLCs acquired a functional hepatic phenotype around day 20, this provides a practical window of approximately 5-6 weeks for chronic disease modelling or long-term exposure studies (Figure S3D). Notably, several important genes involved in hepatic drug metabolism activity such as *CYP3A4*, *CYP2C9* and *CYP2C19* reached their peak expression at day 40 of differentiation. Therefore, while day 20 OBs may already be suitable for applications such as disease modelling, extending differentiation to day 40 is advantageous when the goal is to maximize drug-metabolizing capacity for screening applications. Furthermore, since the SAP system allows co-culture of additional cell types, incorporating non-parenchymal cells alongside hepatocytes could enable the generation of more organotypic models with advanced hepatic functions.

We also demonstrated that the OB model is compatible with automated culture platforms. Although this has not yet been shown for HLCs, we provide proof of principle using HepG2 cells cultured within SAP droplets (Figure S7). The cells remained viable and retained their proliferative capacity for several weeks after encapsulation, while continuing to express key hepatocyte-associated markers, including AFP, ALB, and CK19. Furthermore, HepG2 cells were successfully maintained in SAP droplets using a 384-well plate format with a fully automated liquid handling system for 19–20 days, without loss of a single OB during medium exchanges. These results demonstrate that the OB culture system is highly compatible with automation, an essential feature for high-throughput and large-scale applications.

PuraMatrix^TM^, as a commercial product, is not approved for clinical use, however other RADA16 based products have already paved their way to clinical application. For instance, PuraStat® and PuraBond® both based on RADA16 are already approved for clinical use for wound healing, surgical haemostasis, and endoscopy ^83^. Therefore, these advancements open new avenues for exploring the potential clinical applications of OBs.

Although our model shows strong potential for application in adverse drug reaction assessment (Supplementary Table 3), it still requires further optimization. Despite the high expression of many hepatocyte-specific markers and functions, the levels of several drug-metabolizing enzymes, including CYP1A2, CYP2D6, and CYP2B6, remain lower than those observed in PHHs. While CYP3A4 and CYP2C9 together account for the metabolism of most drugs^84,85^, additional refinements are needed to maximize the predictive power of OBs for drug toxicity screening. Such improvements could involve activation of other, as yet to be identified, transcription factors through genetic engineering or the addition of novel small molecules. Incorporating non-parenchymal cell types may also further enhance metabolic capacity and promote HLC maturation. Moreover, future studies should assess compound diffusion rates and determine whether the peripheral monolayer influences compound transport into and out of OBs.

In conclusion, we present a multi-faceted approach that integrates a simple and defined 3D culture system, an engineered cell line, and an optimized medium to address current challenges in generating scalable, robust, and mature 3D hepatic cultures suitable for long-term and automated applications. Our results demonstrate that this novel strategy enables the formation of organotypic liver-like constructs with advanced hepatic functions, compatible with scale-up and automation for drug toxicity screening. Finally, our findings open promising avenues for developing liver organoid– based protocols with potential clinical applications.

## Acknowledgements

This work was supported by the project RISK-HUNT3R (RISK assessment of chemicals integrating HUman centric Next generation Testing strategies promoting the 3Rs). RISK-HUNT3R has received funding from the European Union’s Horizon 2020 research and innovation programme under grant agreement No 964537 and is part of the ASPIS cluster. This work reflects only the authors’ views, and the European Commission is not responsible for any use that may be made of the information it contains. CV was funded by the Research Foundation, Flanders, Belgium (FWO-S001121N, iPSC-LIMIC) and ZW15-07; DOSOP nr: 3299 Project number: AH.2016.133, request number: 66010, NextGenQBio. MK was supported by Postdoctoral mobility funding from Finnish Cultural Foundation. BT was funded by the Research Foundation, Flanders, Belgium (FWO-1280923N). The iPSC work done at Tampere University was funded by the Research Council of Finland (grant # 353175). Special thanks to Yashwanth Goud Mogulagoni, Sara Van Eyck, Margalida Campaner Socias, Vera Rosado Lobito, and Niels Vidal for their technical assistance. The Computational Biophysics and Imaging Group lead by Jari Hyttinen at Tampere University is acknowledged for their support for creating OPT images. Tampere University iPSC Core and VIB Bioimaging Core Leuven are acknowledged for their services.

## Authors contribution

M.K. designed, planned and executed the study, generated and analysed the data, and wrote the manuscript. S.M. helped with analysis of the bulk RNA sequencing data. B.T. contributed to performing the toxicology assays on 2D HLCs and PHH microtissues. R.M. helped with optimizing the automated liquid handling. G.G. and J.V.C contributed to conduct the CYP450-dependant drug metabolism experiment and its analysis. BB assisted with generating the OPT images. MN provided the fresh PHHs. W.M. provided the 3D PHH microtissues as well as scientific input. K.A. and C.M.V. provided funding for the project, aided with design of the studies and data interpretation. All authors read and edited the manuscript.

