## Supplemental information for "Organobodies: A robust and size-controllable system for generating scalable hiPSC-derived liver organoids for drug toxicity screening"

### Supplementary Figures

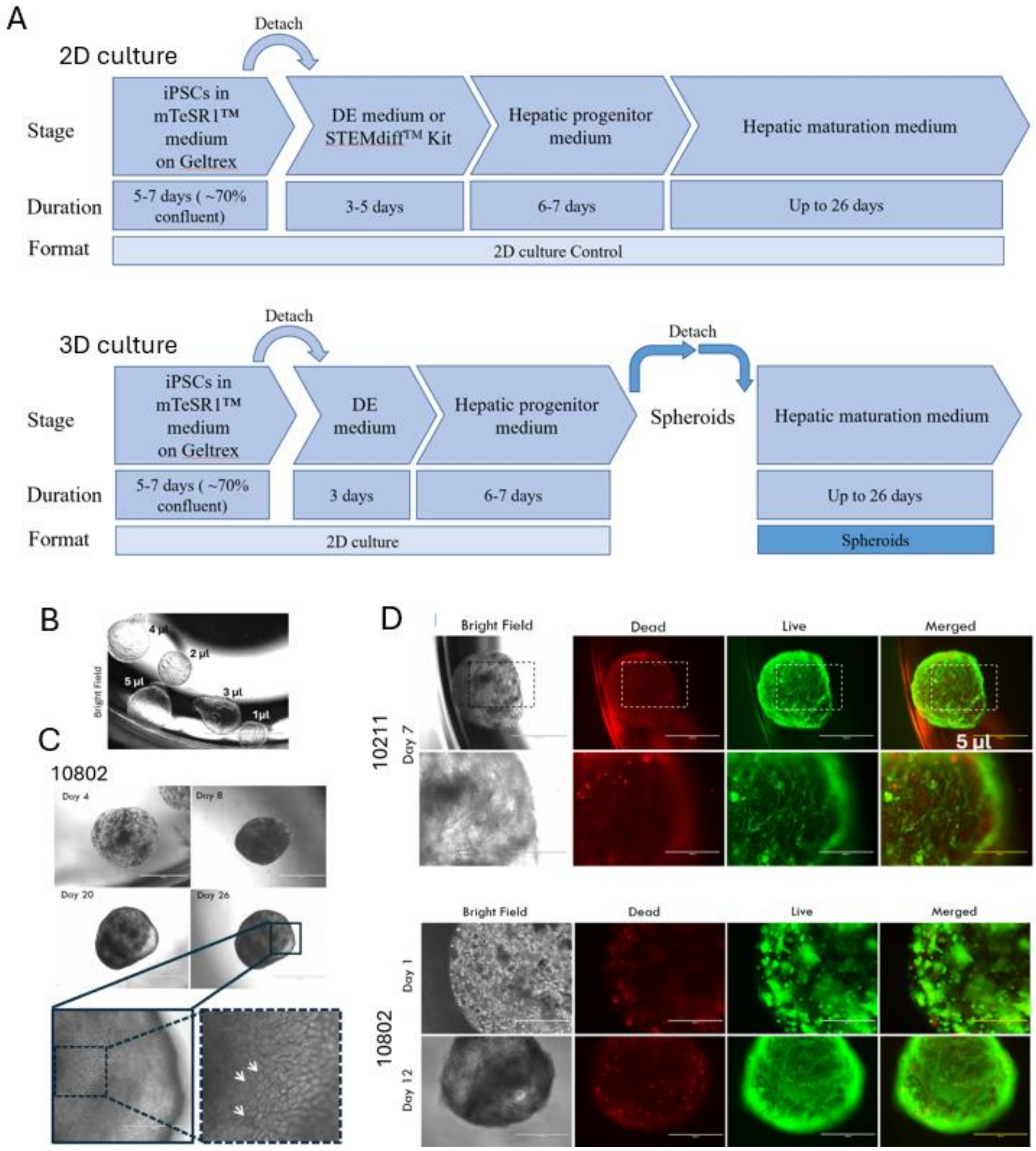


**Figure S1. Generation and characterization of hepatic organobodies (OBs) from cell lines 10211 and 10802. A)** Schematic overview of the differentiation process of 2D hepatocyte-like cells (HLCs) (top panel), and the workflow for generating SAP-cell mix droplets from hepatic progenitors to create hepatic OBs (lower panel). **B)** Brightfield images of SAP hydrogel droplets without cells, generated at varying volumes (1, 2, 3, 4, and 5 µL), imaged using a stereo microscope. **C)** Morphological progression of hepatic OBs derived from cell line 10802, imaged by stereo microscopy on days 4, 8, 20, and 26 post-encapsulation. Lower panels show higher magnification images highlighting the polygonal morphology of hepatocytes at the OB periphery (white arrowheads). **D)** Live/dead staining of OBs derived from lines 10211 (day 7 post-encapsulation) and 10802 (days 1 and 12 post-encapsulation) showing high cell viability within the SAP hydrogel. All droplets were formed using an initial SAP-cell mix volume of 5 µL. Images are representative of one biological replicate with at least five technical replicates per cell line.

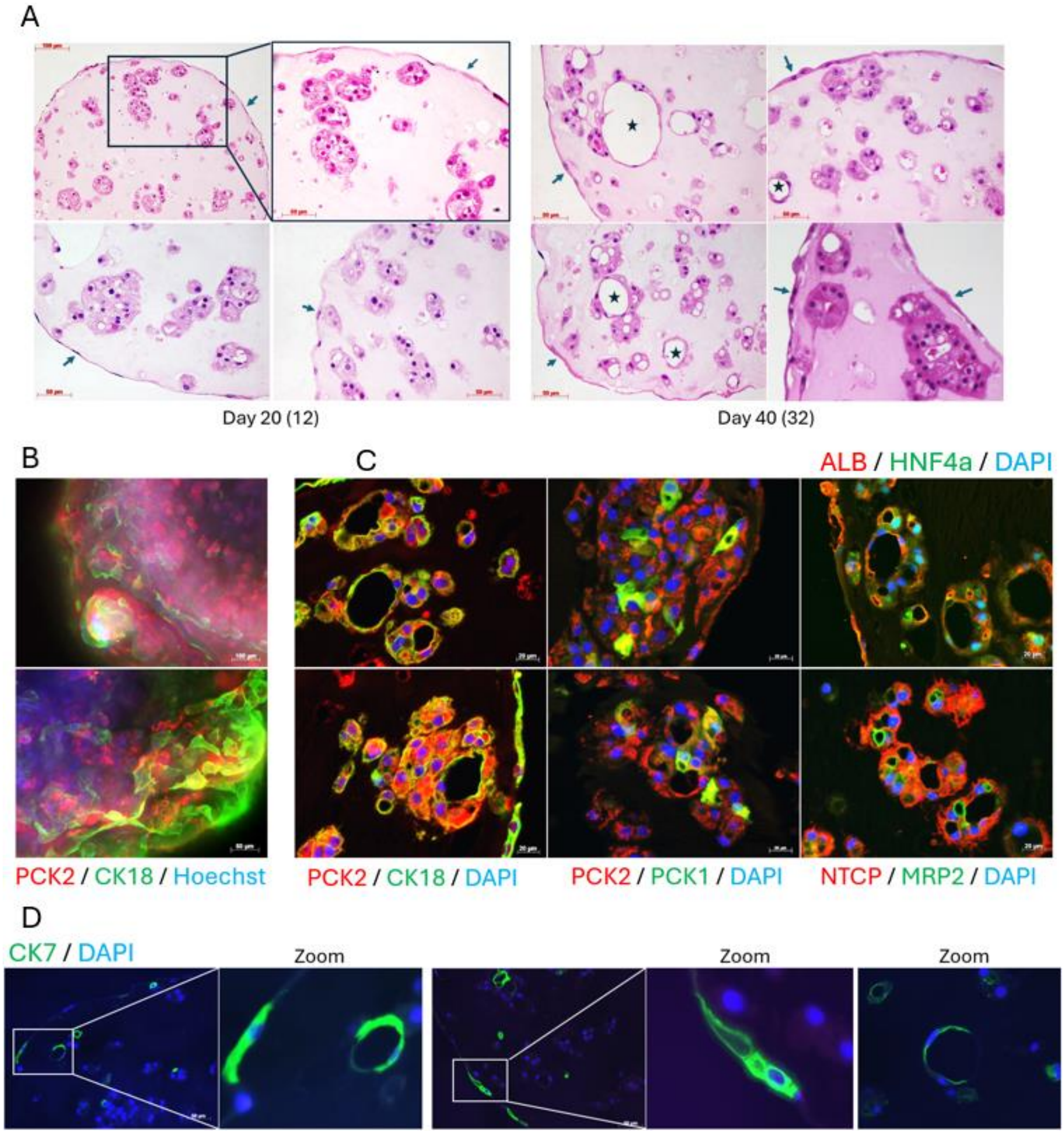


**Figure S2. Histological and immunostaining analysis of low-density hepatic OBs derived from HC3X line. A)** Haematoxylin and eosin (H&E) staining of low-density hepatic OBs on days 20 and 40 of differentiation, corresponding to 12 and 32 days in 3D culture after encapsulation in SAP droplets, respectively (values in brackets). Images show a monolayer of HLCs at the periphery (arrows), with clusters of HLCs located in subperipheral and central regions of the droplets. In prolonged cultures, hepatocytes form lumen-like structures (asterisks). **B)** Whole-mount immunofluorescence staining of low-density HC3X hepatic OBs for the hepatocyte-specific markers PCK2 and CK18. Nuclei were counterstained with Hoechst. **C)** Immunohistochemistry (IHC) staining of OB sections at day 40 (32 days post-encapsulation) showing positive expression of hepatocyte-specific markers PCK2, CK18, PCK1, ALB, HNF4α, NTCP, and MRP2. **D)** IHC staining of OBs for the cholangiocyte marker CK7 at day 40 (32). All images are representative of three independent experiments, each with a minimum of three technical replicates.

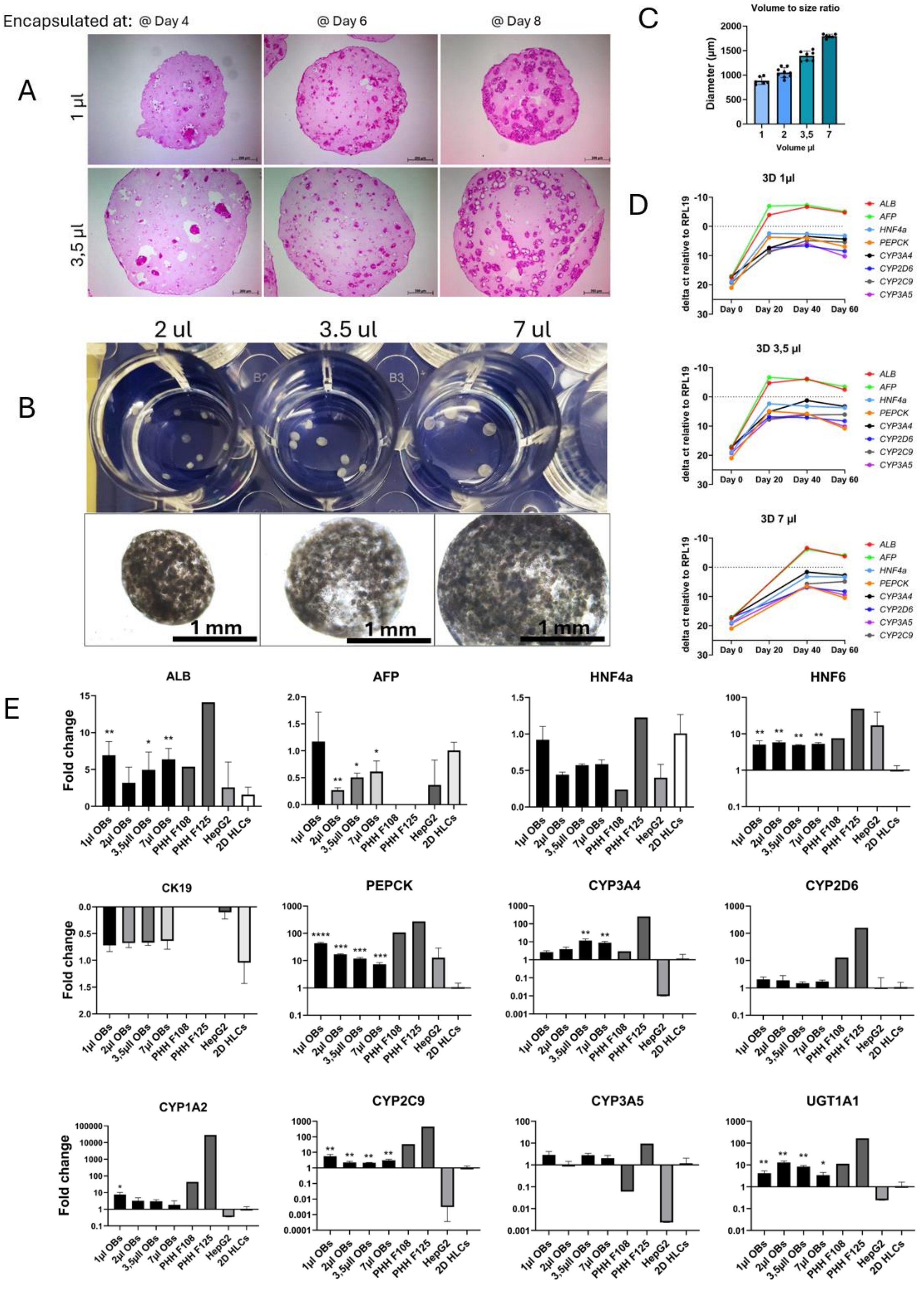


**Figure S3. Optimization and size-dependent analysis of hepatic OBs derived from HC3X line. A)** H&E staining of hepatic OBs generated with cells detached at three distinct stages of differentiation: day 4 (definitive endoderm), day 6 (hepatic commitment), and day 8 (hepatic progenitor stage). OBs were collected on day 40 of differentiation, corresponding to 36, 34, and 32 days post-encapsulation, respectively. **B)** Morphological comparison of OBs generated using different initial volumes (2, 3.5, and 7 μL) of SAP-cell mix. Images were acquired using a digital camera (top panel) and brightfield microscopy (bottom panel). **C)** Quantitative analysis of the volume-to-size relationship of SAP-cell mix droplets, shown in micrometres. **D)** RT-qPCR analysis of hepatocyte-specific gene expression (*ALB*, *AFP*, *HNF4α*, *PEPCK*, *CYP3A4*, *CYP2D6*, *CYP2C9*, and *CYP3A5*) at day 20, day 40, and day 60 of culture. OBs were generated from day 8 hepatic progenitors using three initial SAP-cell mix volumes (1, 3.5, and 7 μL). Data were normalized to RPL19 expression. Each time point includes 3-4 technical replicates. **E)** RT-qPCR comparison of hepatocyte-specific markers (*ALB*, *AFP*, *HNF4α*, *HNF6*, *CK19*, *PEPCK*, *CYP3A4*, *CYP2D6*, *CYP1A2*, *CYP2C9*, *CYP3A5*, *UGT1A1*) in OBs of different sizes (1, 2, 3.5, and 7 μL) harvested on days 37–40. Expression levels were compared with 2D HLCs (day 40), HepG2 cells (2D), and freshly isolated primary human hepatocytes (PHHs; batches F108 and F125). Data are presented as fold changes relative to 2D HLCs. Comparisons were performed using unpaired Student's t-test with Welch's correction for unequal variances. Panels represent data from one biological replicate and 3-5 technical replicates per condition. $P < 0.05$ was considered statistically significant. *$P < 0.05$; **$P < 0.01$; ***$P < 0.001$; ****$P < 0.0001$.

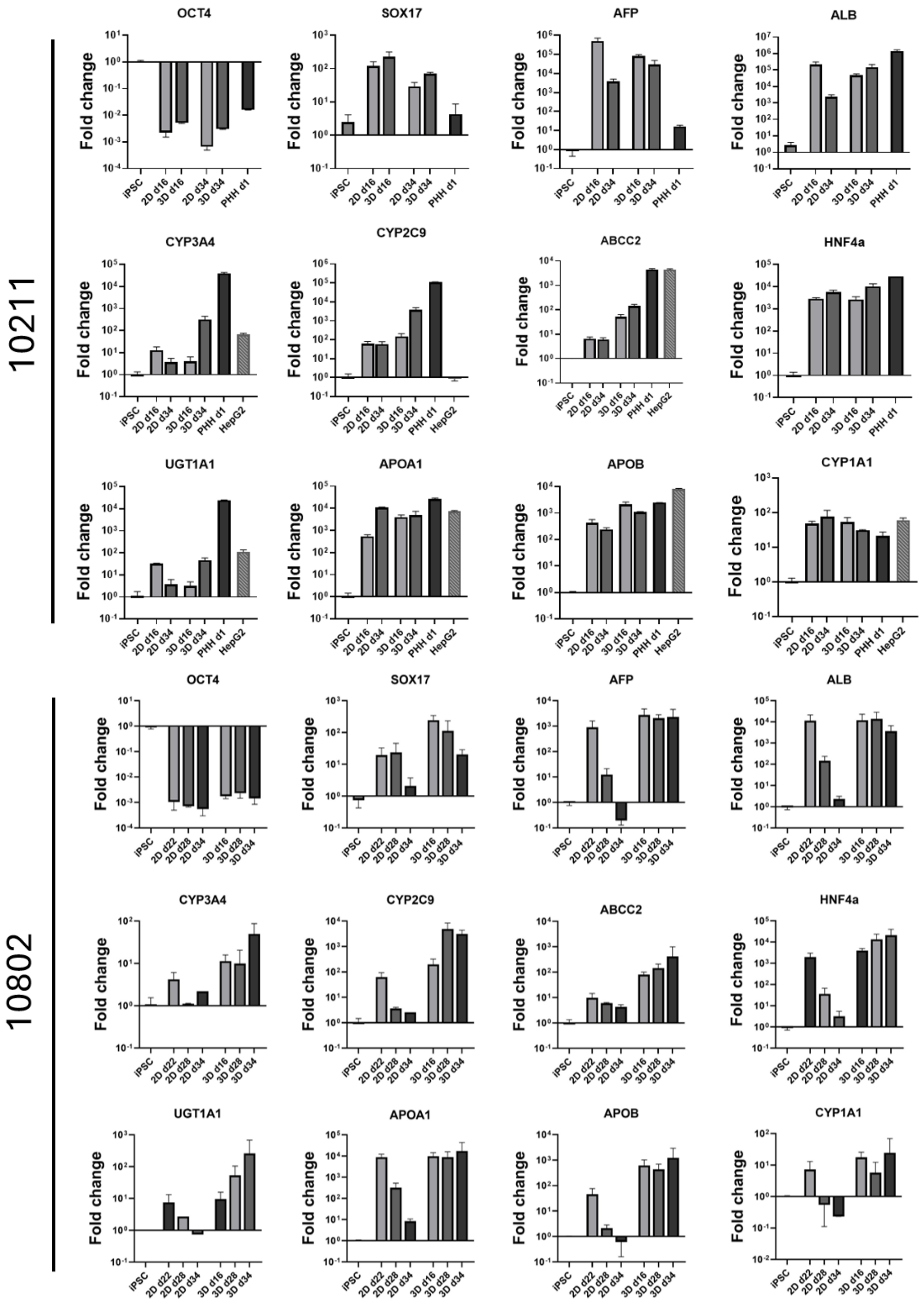


**Figure S4. Comparative gene expression analysis of 2D HLCs and 3D hepatic OBs derived from hiPSC lines 10211 and 10802.** Top panel (cell line 10211): RT-qPCR analysis comparing 2D HLCs and 3D hepatic OBs at days 16 and 34 of differentiation with PHHs (24 h culture) and HepG2 cells (5-day culture). Lower panel (cell line 10802): RT-qPCR analysis of 2D HLCs at days 22, 28, and 34 and corresponding OBs at days 16, 28, and 34. All data were normalized to *GAPDH*, and fold changes were calculated relative to undifferentiated hiPSCs. Data represent one biological replicate with 3–4 technical replicates per condition.

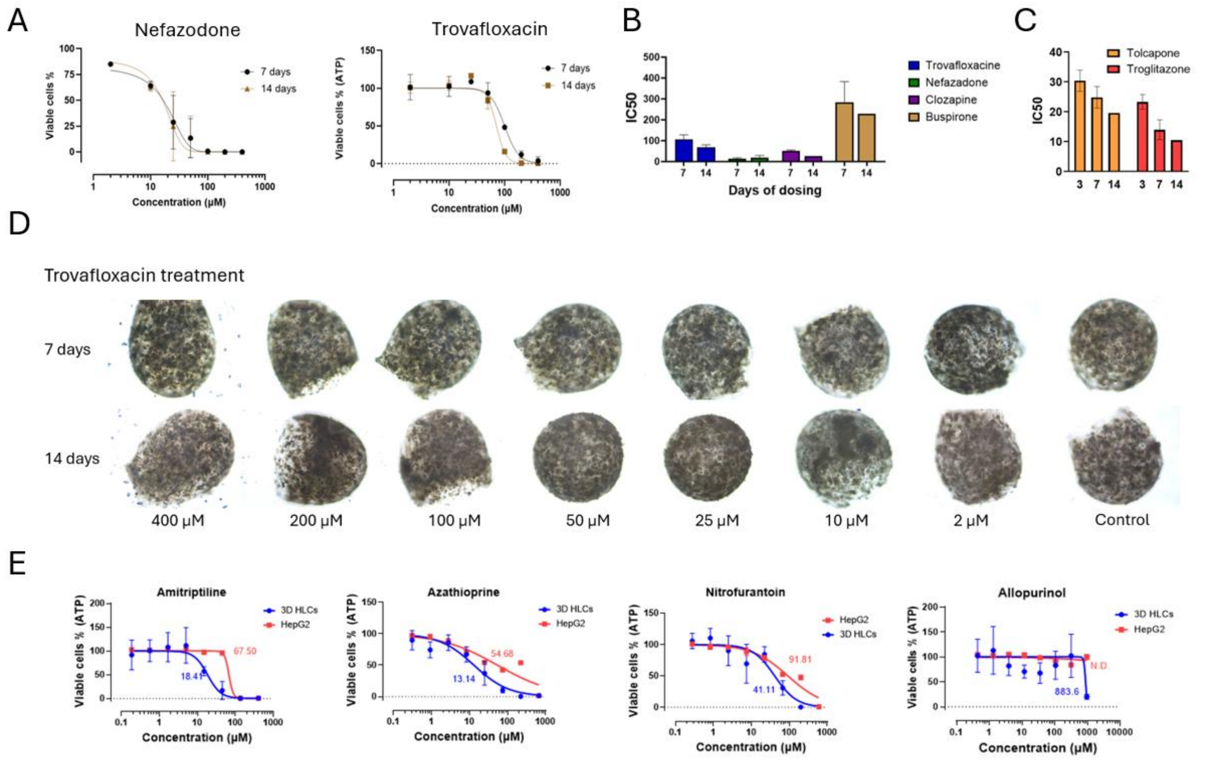


**Figure S5. Extended hepatotoxicity profiling of 3D hepatic OBs. A)** Comparative dose-response toxicity curves of 3D hepatic OBs treated with Nefazodone and Trovafloxacin for 7 and 14 days. Data represent two biological replicates, with a minimum of three technical replicates per dose per replicate. **B)** $IC_{50}$ values calculated from dose–response curves for Trovafloxacin, Nefazodone, Clozapine, and Buspirone after 7- and 14-day treatments. **C)** $IC_{50}$ values for Tolcapone and Troglitazone after 3, 7, and 14 days of treatment, showing increased sensitivity with prolonged exposure. **D)** Morphological assessment of 3D hepatic OBs following Trovafloxacin treatment for 7 and 14 days at various doses. **E)** Comparative dose-response toxicity curves for 3D hepatic OBs and 2D HepG2 cells treated with Amitriptyline, Azathioprine, Nitrofurantoin, and Allopurinol for 7 days. Data represent in this panel is from one biological replicate with a minimum of three technical replicates per dose. $IC_{50}$ values were determined by nonlinear regression analysis of dose–response curves using GraphPad Prism.

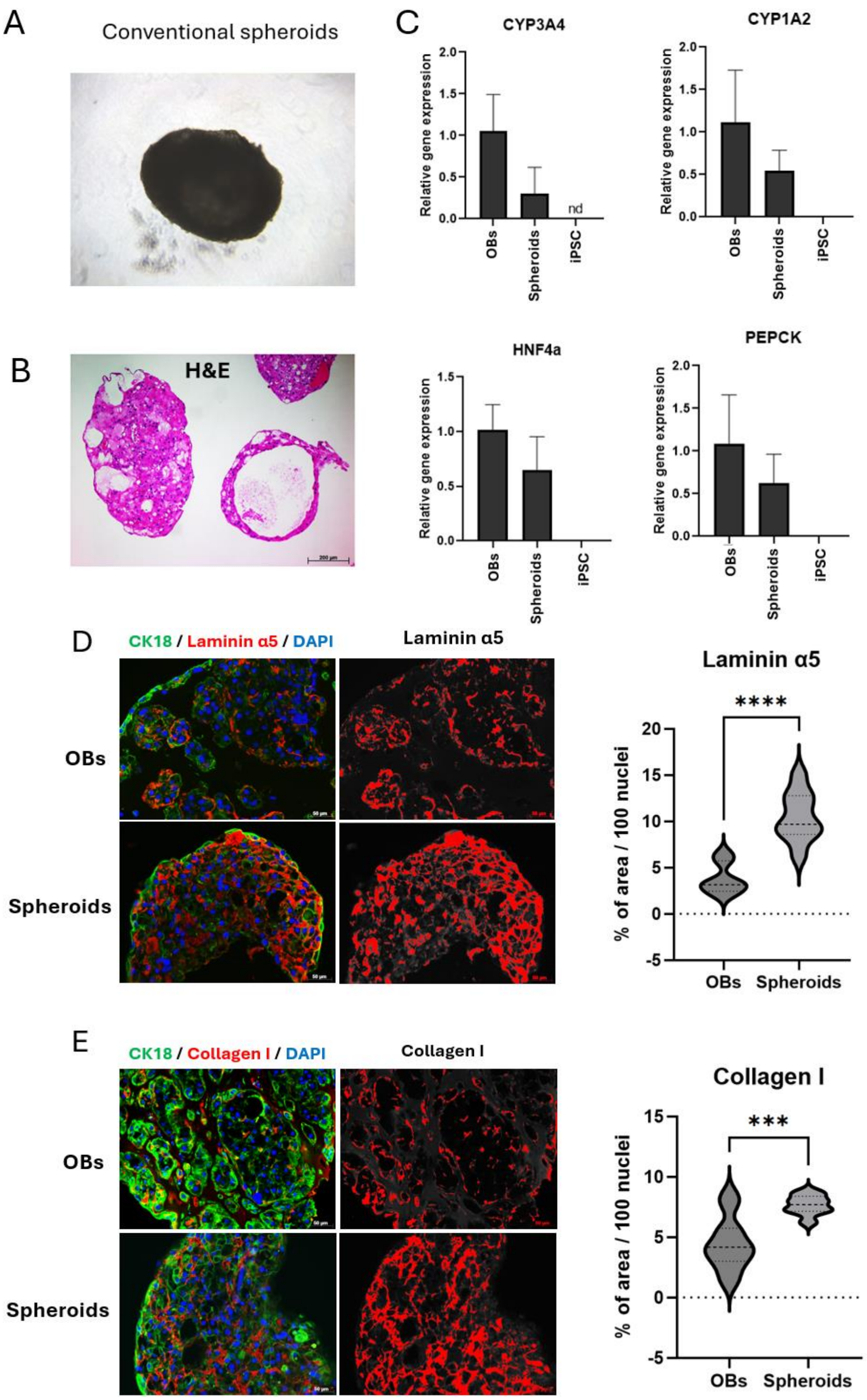


**Figure S6. Comparison of OBs with conventional spheroids both generated from HC3X line. A)** Brightfield and **B)** H&E staining of conventional spheroids generated by self-aggregation in ultra-low attachment plates. **C)** Comparative RT-qPCR analysis of key hepatic genes: *CYP3A4*, *CYP1A2*, *HNF4α*, and *PEPCK*. Quantification of the deposition of **D)** laminin α5 and **E)** collagen I in both OBs and conventional spheroids based on immunohistochemistry images. A collection of 7–11 images were quantified by ImageJ for each analysis and data was normalised based on the nuclei count in each image. The graphs were generated using Graphpad Prism. Comparisons between groups were performed using an unpaired Student's *t*-test with Welch's correction to account for unequal variances. SBs: 3 biological replicates; spheroids: 1 biological replicate. $P < 0.05$ was considered statistically significant. ***$P$ < 0.001; ****$P$ < 0.0001.

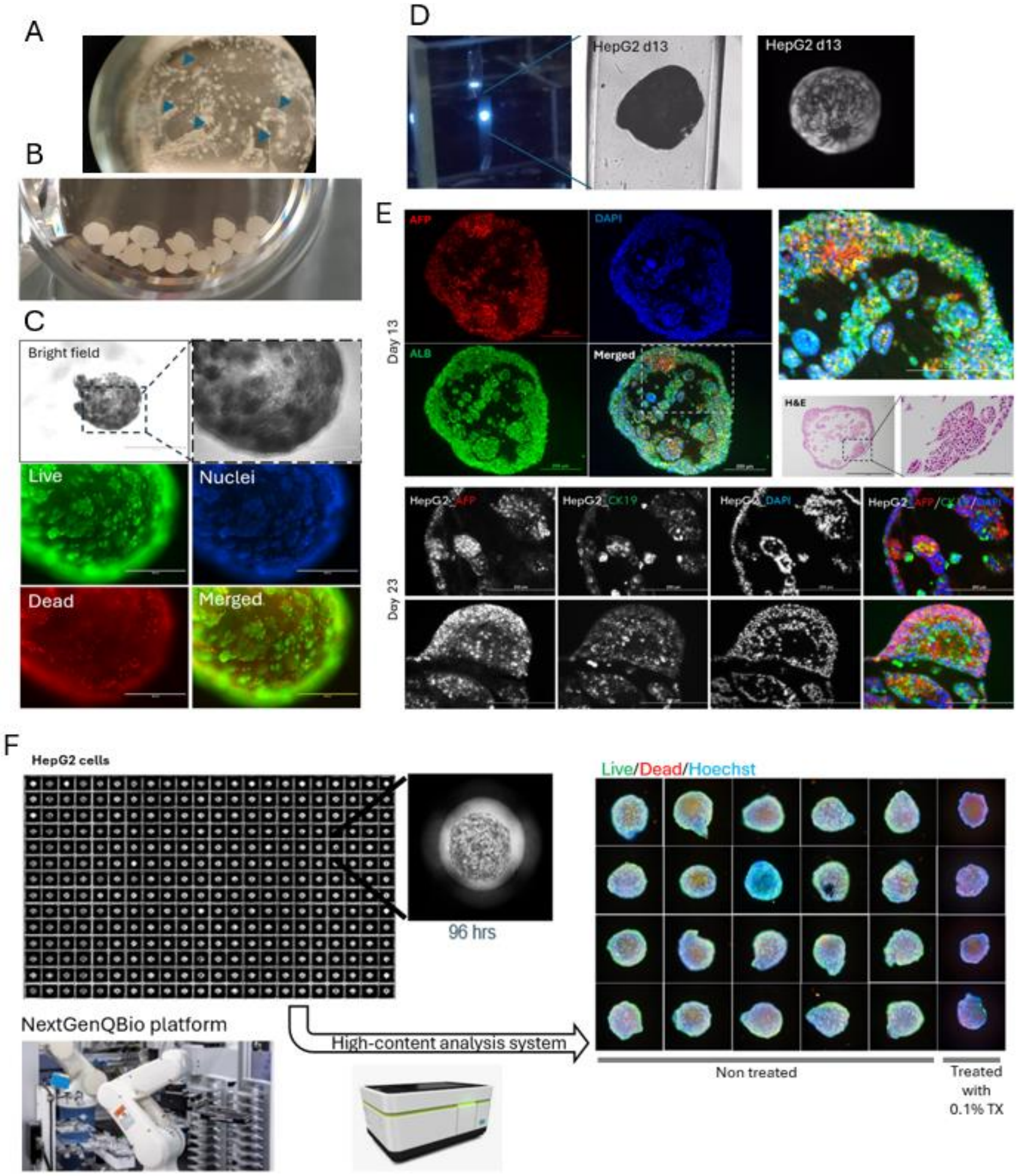


**Figure S7. 3D culture of HepG2 cells in SAP hydrogel. A)** Stereomicroscope image of a 3D HepG2 culture generated using the conventional method, where the SAP-cell mixture is deposited at the bottom of the well and overlaid with medium. This approach results in uneven gelation and partial hydrogel detachment during medium changes (indicated by blue arrows). **B)** Image showing bulk production of multiple HepG2 SAP droplets within a single well. **C)** Brightfield and live/dead staining images of 3D HepG2 SAP droplets at day 13 post-encapsulation. **D)** Optical projection tomography (OPT) image of a 3D HepG2 SAP droplet 13 days after encapsulation. **E)** IHC and H&E staining of HepG2 SAP droplets at days 13 and 23 post-encapsulation, showing positive expression of hepatocyte-specific markers ALB, AFP, and CK19. **F)** High-throughput production of HepG2 SAP droplets in 384-well plates (left panel), maintained under automated liquid handling for 19-20 days. Right panel shows representative wells from a single assay plate following live/dead staining and imaging using the Operetta high-content imaging system (N = 1 biological replicate).

**Supplementary table 1. List of primers used in RT-qPCR study**

| Primer | Forward | Reverse |
|---|---|---|
| *RPL19* | ATTGGTCTCATTGGGGTCTAAC | AGTATGCTCAGGCTTCAGAAGA |
| *AFP* | TGAGCACTGTTGCAGAGGAG | GTGGTCAGTTTGCAGCATTC |
| *ALB* | ATGCTGAGGCAAAGGATGTC | AGCAGCAGCACGACAGAGTA |
| *HNF4α* | ACTACGGTGCCTCGAGCTGT | GGCACTGGTTCCTCTTGTCT |
| *NTCP* | ATCGTCCTCAAATCCAAACG | CCACATTGATGGCAGAGAGA |
| *CYP3A4* | TTCCTCCCTGAAAGATTCAGC | GTTGAAGAAGTCCTCCTAAGCT |
| *CYP2D6* | CATACCTGCCTCACTACCAAA | TGTCCTGCCTGGTCCTC |
| *G6PC* | GTGTCCGTGATCGCAGACC | GACGAGGTTGAGCCAGTCTC |
| *PEPCK* | AAGAAGTGCTTTGCTCTCAG | CCTTAAATGACCTTGTGCGT |
| *CYP2C9* | GTTTCTGCCAATCACACGTTC | CTGCAGTTGACTTGTTTGGAG |
| *CYP2C19* | CAATGATAGTGGGAAAATTATTGCAT | GTGATCTGCTCCATTATTTCCA |
| *CYP3A5* | GCCCTGAAAGGTTCAGT | GTTGAAGAAGTCCTTGCG |
| *HNF6* | AAATCACCATTTCCCAGCAG | ACTCCTCCTTCTTGCGTTCA |
| *CK19* | CGACTACAGCCACTACTACAC | GGTGGCACCAAGAATCTTGTC |
| *CYP1A2* | CAGCTCTGGGTCATGGTTG | CCTCCTTCTTGCCCTTCAC |
| *UGT1A1* | CAACTGCCTTCACCAAAATCCA | GCAAGATTCGATGGTCGGGTT |

**Supplementary Table 2. List of antibodies**

| Type | Antibody | Cat. No | Company | Host | Dilution | Application |
|---|---|---|---|---|---|---|
| **Primary antibodies** | HNF4α | ab41898 | abcam | Mouse | 1:200 | IHC |
| | NTCP | HPA042727 | Sigma-Aldrich | Rabbit | 1:250 | IHC |
| | α1-Antitrypsin | A0012 | Dako | Rabbit | 1:200 | IHC |
| | CYP3A4 | Bs90368 | BioWorld Technology | Rabbit | 1:100 | IHC/ICC |
| | HNF6 | Ab186743 | abcam | Rabbit | 1:200 | IHC |
| | PCK1 | sc-377027 | Santa Cruz | Mouse | 1:200 | IHC |
| | PCK2 | Sc-32879 | Santa Cruz | Rabbit | 1:200 | IHC/ICC |
| | Albumin | A0001 | Dako | Rabbit | 1:200 | IHC |
| | CK18 | M7010 | Dako | Mouse | 1:200 | IHC/ICC |
| | MRP2 | Ab3373 | abcam | Mouse | 1:200 | IHC |
| | CK7 | MAB3226 | Chemicon International | Mouse | 1:200 | IHC |
| **Secondary antibodies** | **Antibody** | **Cat. No** | **Company** | **Host** | **Dilution** | **Application** |
| | anti-Rabbit IgG (H+L) Alexa Fluor 647 | A-31573 | Thermo Fisher Scientific | Donkey | 1:500 | IHC/ICC |
| | anti-Sheep IgG (H+L) Alexa Fluor 488 | A-11015 | Thermo Fisher Scientific | Donkey | 1:500 | IHC/ICC |
| | anti-Rabbit IgG (H+L) Alexa Fluor 555 | A-31572 | Thermo Fisher Scientific | Donkey | 1:500 | IHC/ICC |
| | anti-Mouse IgG (H+L) Alexa Fluor 647 | A-31571 | Thermo Fisher Scientific | Donkey | 1:500 | IHC/ICC |
| | anti-Mouse IgG (H+L) Alexa Fluor 555 | A-31570 | Thermo Fisher Scientific | Donkey | 1:500 | IHC/ICC |
| | anti-Rabbit IgG (H+L) Alexa Fluor 488 | A-21206 | Thermo Fisher Scientific | Donkey | 1:500 | IHC/ICC |
| | anti-Mouse IgG (H+L) Alexa Fluor 488 | A-21202 | Thermo Fisher Scientific | Donkey | 1:500 | IHC/ICC |

**Supplementary Table 3.** Comparative table providing calculated $IC_{50}$ values for compounds already tested available in published studies.

| **Compound** | **DILI concern**** | **Kiamehr et al.** | **Qosa et al. (Qosa *et al.*, 2021)** | **Kim et al. (Kim *et al.*, 2022)** | **Lee et al. (Lee *et al.*, 2021)** | **Proctor et al. (Proctor *et al.*, 2017)** | **Feng lee (Li *et al.*, 2020)** |
|---|---|---|---|---|---|---|---|
| **Culture set up** | | HLCs in RADA16 OBs | HLCs conventional spheroids in ULA plates | iPSC organoids in Matrigel | 3D HLCs microwells | 3D PHH microtissues from InSphero | 3D PHH spheroids in ULA |
| **Days of exposure** | | 7 days | 72 hrs | 1 day | 7 days | 14 days | 5 days |
| **Trovafloxacin** | 8 | 111,8 | | | | >125 | 55 |
| **Clozapine** | 5 | 50,18 | 43,2 | | | 34,8 | 33,6 |
| **Nefazodone** | 8 | 10,74 | | ~106 | 86,11 | 29,4 | 5,2 |
| **Buspirone** | 3 | 291,1 | | NT* | | 163 | |
| **Acetaminophen** | 5 | 7354 | 22400 | ≈33880 | 29370 | 572 | 927 |
| **Obeticholic acid** | - | 25,87 | | | | | |
| **Tolcapone** | 8 | 26,23 | | | | 19 | 19,3 |
| **Entacapone** | 0 | 77,44 | | | | 152,3 | 125,1 |
| **Troglitazone** | 8 | 14,26 | 45,2 | ≈204 | 133,4 | 14,6 | 1 |
| **Amitriptyline** | 5 | 18,4 | | | | 18,8 | 9,7 |
| **Azathioprine** | 5 | 13,14 | | | | 22,2 | <7,8 |
| **Nitrofurantoin** | 8 | 41,11 | | | | 69,1 | 17 |
| **Allopurinol** | 8 | 883,6 | | | | | |

*: NT = no toxicity was detected

**: 0 = least DILI concern; 8 = most DILI concern
